# Dynamics of synthetic transcriptional condensates emerge from RNA synthesis and degradation

**DOI:** 10.64898/2026.05.03.722550

**Authors:** Jane Liao, So Yeon Ahn, Allie C. Obermeyer

## Abstract

At sites of active gene expression, dynamic compartments known as transcriptional condensates assemble and dissolve on timescales relevant to RNA synthesis and degradation. Yet how the non-equilibrium dynamics of these condensates emerge from the coupling of RNA concentration and phase separation remains poorly understood. Here we engineer synthetic active condensates in which T7 RNA polymerase transcribes RNA in situ, triggering phase separation with a cationic scaffold protein. By using RNA concentration as a tunable parameter, we drive condensates along defined paths through a characterized phase diagram. This reaction-phase separation coupling gives rise to three emergent dynamic phenomena not accessible in passive systems: a rapid switch-like nucleation burst, RNA-mediated positive and negative feedback regulation of transcription, and oscillatory condensate formation in which RNA degradation spontaneously renucleates condensates. Together, these results show that the dynamic functions of transcriptional condensates emerge from their reaction-driven paths through phase space, providing a quantitative framework for understanding how RNA flux governs condensate dynamics in living cells.

## Introduction

Inside the crowded intracellular environment, cells regulate gene expression with remarkable spatial and temporal precision. Increasing evidence suggests that this regulation is facilitated in part by biomolecular condensates that organize the transcription machinery into dynamic mesoscale compartments [1, 2]. Unlike membrane-bound organelles or stoichiometric complexes, transcriptional condensates arise through multivalent interactions that concentrate proteins and nucleic acids while remaining highly dynamic, assembling, dissolving, and reorganizing on timescales comparable to RNA synthesis [3]. Such responsiveness suggests that transcriptional condensates are not merely passive compartments, but active reaction-coupled assemblies with compositions and lifetimes that evolve with gene expression.

While substantial progress has been made in identifying the molecular constituents and interaction motifs of transcriptional condensates, a central question remains about how time-dependent functions emerge from the coupling between RNA concentration and phase separation. In cells, RNA is continuously produced by transcription and degraded by ribonucleases, creating a dynamic flux that can reshape condensate composition and material properties over time [4–6]. Yet most in vitro studies reconstitute condensates under static, equilibrium conditions, rather than using active biochemical reactions to drive condensates along the relevant kinetic paths [7, 8]. Consequently, the dynamic functions of condensates in living systems remain difficult to reproduce in vitro and probe mechanistically [9, 10].

To provide a systematic framework linking transcriptional activity to condensate dynamics, we create synthetic active transcriptional condensates in a fully defined system, where T7 RNA polymerase synthesizes RNA in situ to trigger associative phase separation with a cationic scaffold protein. Using characterized phase boundaries to predict and navigate reaction-driven paths, we observe rapid nucleation bursts recapitulating switch-like condensate assembly at active genomic loci [11–13], and RNA-mediated feedback regulation of transcription consistent with RNA-driven condensate regulation in living cells [4]. We also show that RNA degradation by ribonuclease drives the reverse path through the phase diagram, and combining sequential RNA synthesis and degradation leads to oscillatory condensate formation, a dynamic behavior not previously reconstituted in a defined biochemical system. Collectively, these results demonstrate that condensate dynamics and biological function emerge from reaction-driven paths through phase space, providing a quantitative framework for understanding how RNA concentration governs transcriptional condensate behavior in cells. Beyond serving as a minimal physical model for transcriptional condensates, this framework provides a foundation for dynamically compartmentalizing genetic information in synthetic systems and programming life-like temporal organization in adaptive protocell-like systems [14–23].

## Results

### Design of synthetic active transcriptional condensates

For transcriptional condensates, RNA concentration is a governing parameter that is difficult to vary systematically in cellular systems, as perturbing RNA levels causes broader physiological consequences. Reconstituting transcriptional condensates in vitro allows direct dissection of how RNA flux drives condensate dynamics, but requires deliberate design choices to access the full range of biologically relevant behaviors. Here we investigated an associative phase separation framework, rather than homotypic nucleic acid self-assembly, to better capture the protein-RNA electrostatic interactions that contribute to endogenous transcriptional condensate formation [24, 25]. A key design constraint is that the scaffold interaction strength must be tuned within a narrow range, strong enough to drive phase separation but weak enough to be reversible [26, 27]. Given this constraint, we engineered GFP(+4), a cationic scaffold with net charge sufficient to drive phase separation with RNA in a defined transcription reaction buffer (Supplementary Fig. 1-3) [28, 29]. Our goal was not to reconstitute the full complexity of endogenous transcriptional condensates, but to isolate RNA concentration as the primary variable driving condensate dynamics. We therefore used a minimal T7 RNAP-based system for precise and tunable control over RNA synthesis.

We first mapped the phase diagram of this system under passive conditions by mixing purified RNA with GFP(+4) in transcription buffer without the polymerase to provide an equilibrium reference for our active system (Fig. 1a). Turbidity measurements confirmed that phase separation required both RNA and GFP(+4), consistent with electrostatically driven associative phase separation (Fig. 1b). Phase separation was confined to an intermediate range of RNA-to-protein ratios centered around charge neutrality and suppressed by excessive charge imbalance at either extreme, delineating a two-phase region with both a lower and a higher RNA boundary (Fig. 1c). The higher RNA boundary, known as a reentrant boundary, arises when excess RNA inverts the charge of the condensate, destabilizing the dense phase [30]. This implied a critical prediction for the active system. As RNA accumulates during transcription, condensates should nucleate, grow, and ultimately dissolve as the system traverses the two-phase region at fixed protein scaffold concentration.

**Fig. 1.**
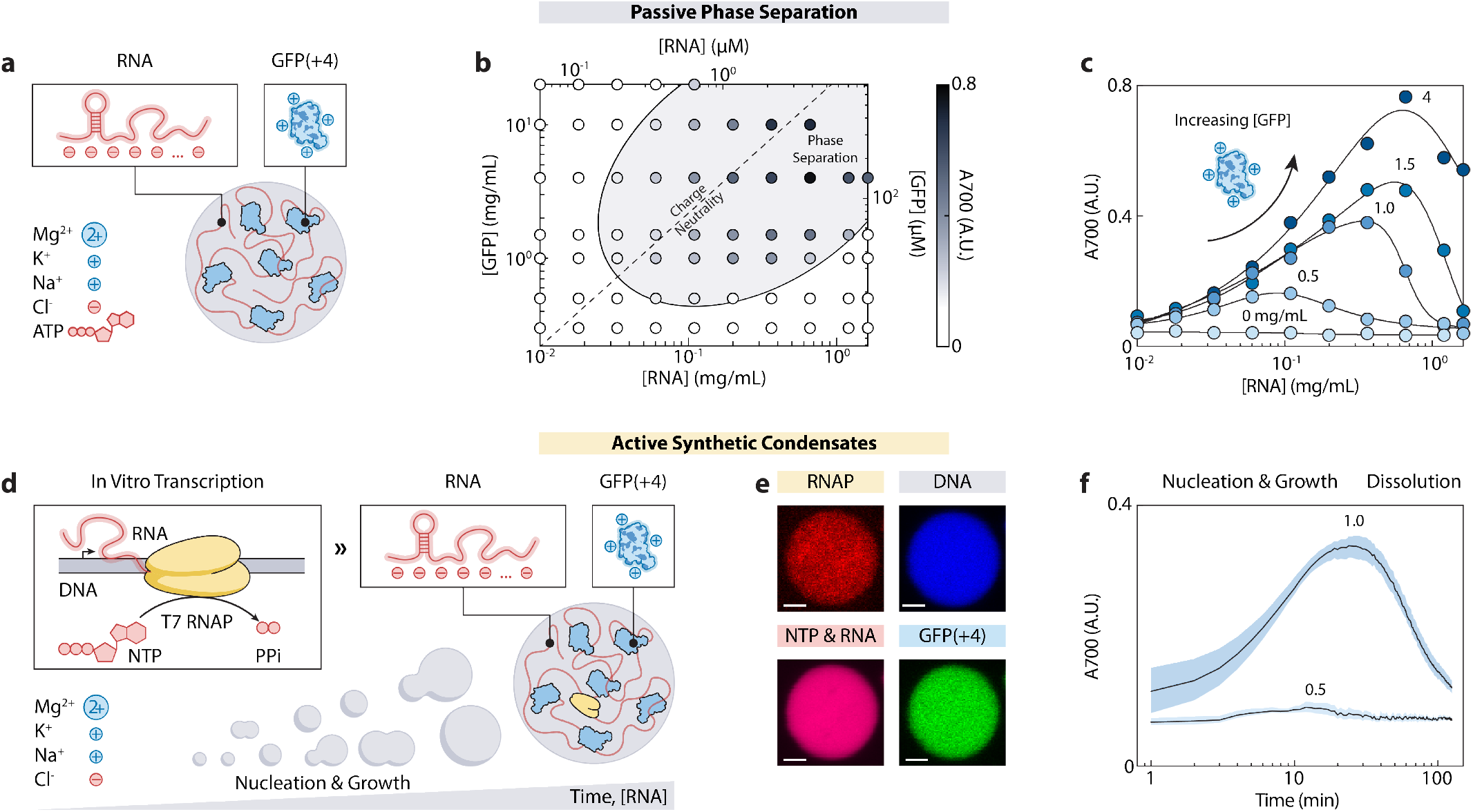
Cationic GFP and RNA phase separate in passive and active systems. **a.** Schematic of passive phase separation in which purified RNA (531 nt, 171 kDa) and GFP(+4) (28 kDa) were mixed in a buffer containing 40 mM HEPES, 80 mM KCl, 8 mM MgCl2, and 6 mM ATP at pH 7.4. **b**. Phase diagram showing the approximate two-phase region (shaded) of passive condensates with dashed line representing charge neutrality assuming an RNA charge density of 1*e* per nucleotide. Marker color represents the absorbance at 700 nm (A700). **c**. Turbidity as a function of RNA concentration at fixed GFP(+4) concentrations (markers), replotted from b. to highlight reentrant dissolution at high RNA-to-protein ratios. **d**. Schematic of synthetic active condensates where T7 RNAP-driven transcription of a linear dsDNA template triggers the in-situ nucleation and growth of synthetic condensates. **e**. Confocal images showing the co-partitioning of T7 RNAP (Alexa Fluor 594), linear template DNA (DAPI), NTP and RNA (Cy5-UTP), and GFP(+4) within an active condensate after 60 min of IVT. GFP(+4) concentration was 1 mg mL^-1^. Scale bars, 2 *µ*m. **f**. Turbidity (A700) over time during IVT-driven condensate nucleation and reentrant dissolution at fixed GFP(+4) concentrations (0.5 and 1 mg mL^-1^). Data represent mean and range of *n* = 3 independent reactions. Reaction conditions for e. and f. were 76 nM linear DNA template, 100 nM T7 RNAP, 40 mM HEPES (pH 7.4), 80 mM KCl, 8 mM MgCl2, and 6 mM total NTP.

To test this, we replaced purified RNA with an in vitro transcription (IVT) reaction in which T7 RNAP synthesizes RNA from a linear dsDNA template in situ (Fig. 1d). Time-course turbidity measurements and confocal microscopy confirmed that condensates nucleated and grew within minutes as transcription proceeded, and fluorescence labeling showed RNA and GFP(+4) co-condensing in active condensates, mirroring the passive system (Fig. 1e, f). Interestingly, both the DNA template and T7 RNAP were also enriched within the active condensates as client molecules (Fig. 1e, Supplementary Fig. 4). As predicted from the passive phase diagram, the active system exhibited non-monotonic turbidity over time with an initial rise as condensates nucleated followed by reentrant dissolution as RNA exceeded the charge-balanced regime, consistent with the active condensates traversing a reaction-driven path through the two-phase region and past the reentrant phase boundary (Fig. 1f).

### Active condensate growth and dissolution can be controlled

The direct coupling between RNA flux and condensate dynamics raised the question of how different condensate behaviors emerge from transcriptional activity. We hypothesized that condensate dynamics are governed by the rate of RNA synthesis and the concentration of RNA-binding scaffold proteins, two parameters that determine how the system traverses the phase diagram. Critically, these parameters can also be regulated by cells through gene expression. We therefore systematically varied these two parameters to map their effects on active condensate behavior.

We expected that higher concentrations of T7 RNAP would increase the rate of RNA production, leading to an earlier onset of nucleation as the system exceeds the saturation concentration. Furthermore, we anticipated that a higher transcription rate would accelerate condensate growth, driven by both diffusion-limited growth and Brownian-mediated coalescence. To quantify this, we performed time-lapse microscopy of IVT reactions using a fixed initial concentration of the GFP(+4) scaffold protein (2.5 mg mL^-1^) and varying concentrations of T7 RNAP (40, 100, 200 nM) (Fig. 2a). Following image segmentation, we quantified the increase in the number-weighted mean radius ⟨*r*⟩*N* over time (Fig. 2b). Consistent with our hypothesis, higher enzyme concentrations produced more RNA, accelerating the nucleation of active condensates (Fig. 2c). Increasing the T7 RNAP concentration from 40 to 200 nM resulted in an earlier nucleation burst and a shorter time to reach peak condensate number density (Supplementary Fig. 5). Furthermore, the growth rate in condensate radius, *d* ⟨*r*⟩*N/dt*, between 20-30 min was linearly correlated with the concentration of T7 RNAP in this range, showing that T7 RNAP concentration is a robust control parameter for both nucleation and subsequent growth of active condensates (Supplementary Fig. 6).

**Fig. 2.**
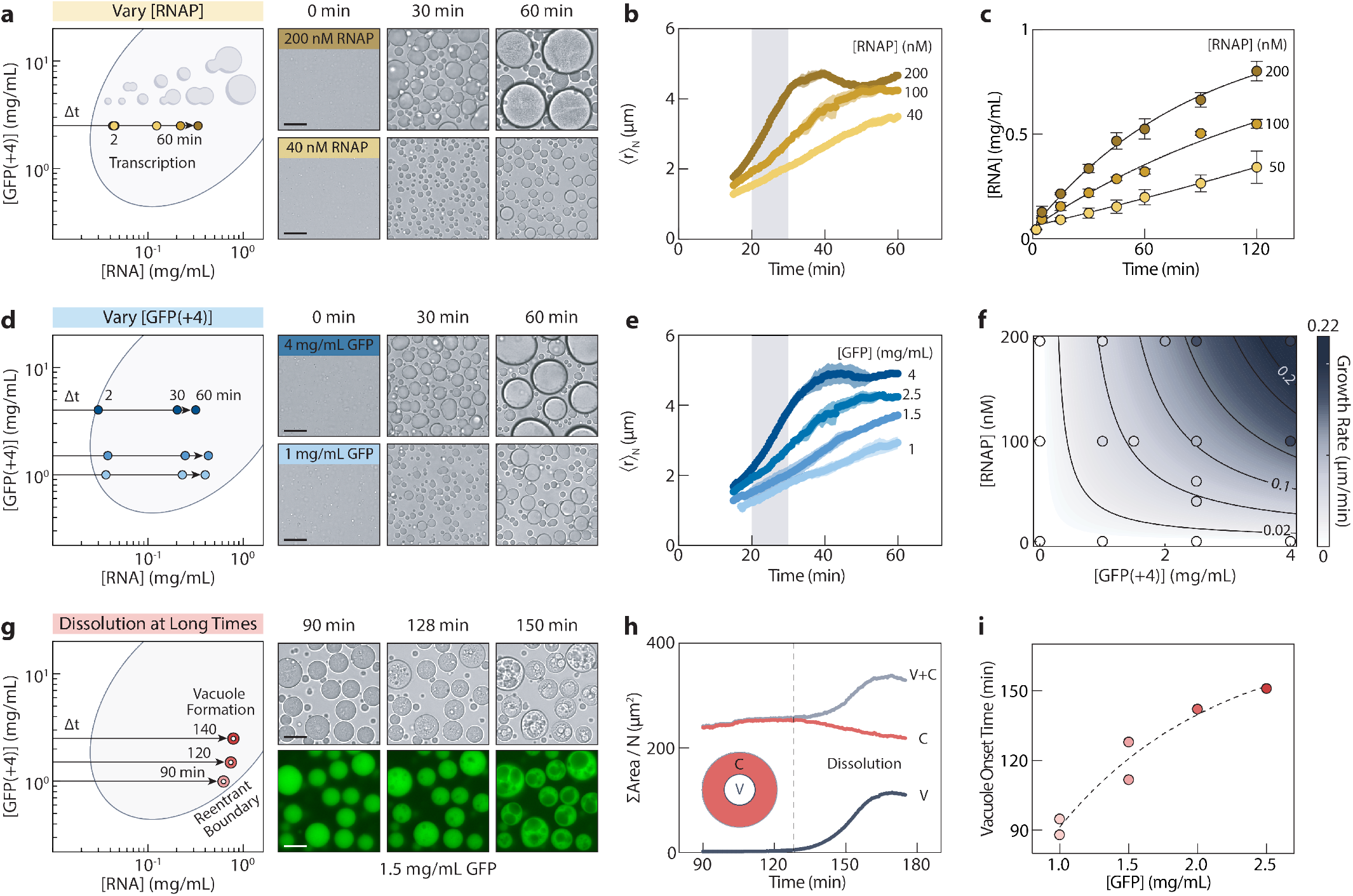
Active condensate growth and dissolution can be controlled by enzyme and cationic scaffold concentrations. **a.** Phase diagram and reaction-driven paths during transcription with representative microscopy images showing the condensate growth over time with varying T7 RNAP concentrations and a fixed concentration of 2.5 mg mL^-1^ GFP(+4). Data points correspond to measured RNA concentrations at the indicated reaction times. Scale bars, 20 *µ*m. **b**. Time-course of mean condensate radii (number-weighted) from bright-field microscopy show that growth kinetics increase with T7 RNAP concentration. Shaded areas denote the range of *n* = 2 independent replicates. The gray region (20–30 min) indicates the window used for growth rate estimations in (f). **c**. Total RNA concentration over time under the conditions corresponding to a and b, as quantified by SYTO 63 fluorescence. Error bars represent the range of 3 independent replicates. **d**. Same as in (a), but with varying concentrations of GFP(+4) scaffold and a fixed concentration of 100 nM T7 RNAP. Scale bars, 20 *µ*m. **e**. Same as in (b), but with varying concentrations of GFP(+4). f. The growth rate of condensates can be controlled by both GFP(+4) and T7 RNAP concentrations. The increase in ⟨*r*⟩ *N* between 20 and 30 min was estimated by linear regression and then fit to a multiplicative saturation model. Black contour lines denote constant predicted growth rates. **g**. Same as in (a), but with 200 nM T7 RNAP and varying GFP(+4) concentrations at long times with representative microscopy images showing vacuole formation over time. Scale bars, 20 *µ*m. **h**. Average vacuole, V, condensate, C, and total area, V+C, normalized by the number of detected condensates in the frame. Dashed vertical line corresponds to the vacuole onset time as defined by a vacuole area fraction threshold of 0.02 by Otsu thresholding of the GFP channel. **i**. Vacuole onset time from image analysis increases with the concentration of GFP(+4). Dashed line represents an exponential plateau fit.

To determine how the cationic scaffold concentration shapes emergent condensate dynamics, we next varied the GFP(+4) concentration while keeping the concentration of T7 RNAP fixed. While the T7 RNAP concentration determines the rate and amount of RNA production, the GFP(+4) concentration sets the amount of cationic scaffold available for associative phase separation with RNA. We found that higher concentrations of the cationic scaffold significantly increased both nucleation and growth rates (Fig. 2d, e). Notably, the number-weighted mean radius after 1 h was strongly dependent on the concentration of the cationic scaffold. This trend likely reflects the geometry of the phase diagram; at lower cationic scaffold concentrations, reaction-driven paths through the phase diagram end closer to the dilute phase boundary after 1 h, resulting in a lower dense phase volume fraction (Supplementary Fig. 5c). Condensate growth kinetics across a range of T7 RNAP and cationic scaffold concentrations revealed that the growth rate increases with both parameters before saturation (Fig. 2f). This behavior is well-described empirically by a multiplicative saturation model, demonstrating that T7 RNAP and cationic scaffold concentration can be used as two independent parameters for tunable control over condensate growth rate (Supplementary Fig. 6).

Extending our analysis beyond the initial nucleation and growth phase, we also characterized the late-stage condensate dynamics as excess accumulating RNA drove the system across the two-phase region toward the reentrant binodal boundary. At high T7 RNAP concentration (200 nM), we observed the formation of hollow internal vacuoles at long times, consistent with vacuole formation during reentrant phase transitions in similar systems [30–33]. Quantitative image analysis revealed that condensates swell as the vacuole void expands, while the area of the dense phase in the rim decreases, indicating net displacement of condensate material and the onset of dissolution as the system approaches the reentrant boundary. The vacuole onset time increased with the initial cationic scaffold concentration, from approximately 90 to 150 min across a range of 1 to 2.5 mg mL^-1^ GFP(+4). This trend reflects the width of the two-phase region, where the reentrant binodal boundary shifts to higher RNA concentrations as the cationic scaffold concentration increases, meaning that more RNA must accumulate before vacuole formation is triggered. Interestingly, nucleolar vacuoles in plant cells are also associated with high transcriptional activity, where an imbalance between ribosomal ribonucleoprotein production and transport creates internal voids [34], suggesting that vacuole formation may be a result of material flux in transcriptionally active condensates. By characterizing RNA synthesis in model transcriptional condensates in vitro, we demonstrate that condensate lifetimes and the onset of remodeling events such as vacuole formation can emerge as physical consequences of transcriptional activity and scaffold protein concentration (Supplementary Fig. 7).

### Active condensates rapidly nucleate in response to transcription

Having mapped the passive phase boundary and characterized non-equilibrium reaction-driven paths through the phase diagram, we next sought to understand the functional consequences of condensate formation. Motivated by observations that RNA Pol II clustering is regulated and correlated with the cell’s ability to mount rapid transcriptional responses, we asked whether analogous switch-like condensate nucleation could arise in our synthetic active system through reaction-phase separation coupling alone [35]. To test this, we performed IVT at high cationic scaffold concentration (10 mg mL^-1^) and low T7 RNAP concentration (50 nM), conditions that drive the system gradually toward charge neutrality as RNA accumulates over the course of an hour. As a passive reference, we mixed GFP(+4) with purified RNA at a concentration approximately equal to the final RNA yield (0.15 mg mL^-1^) from IVT (Fig. 3a).

**Fig. 3.**
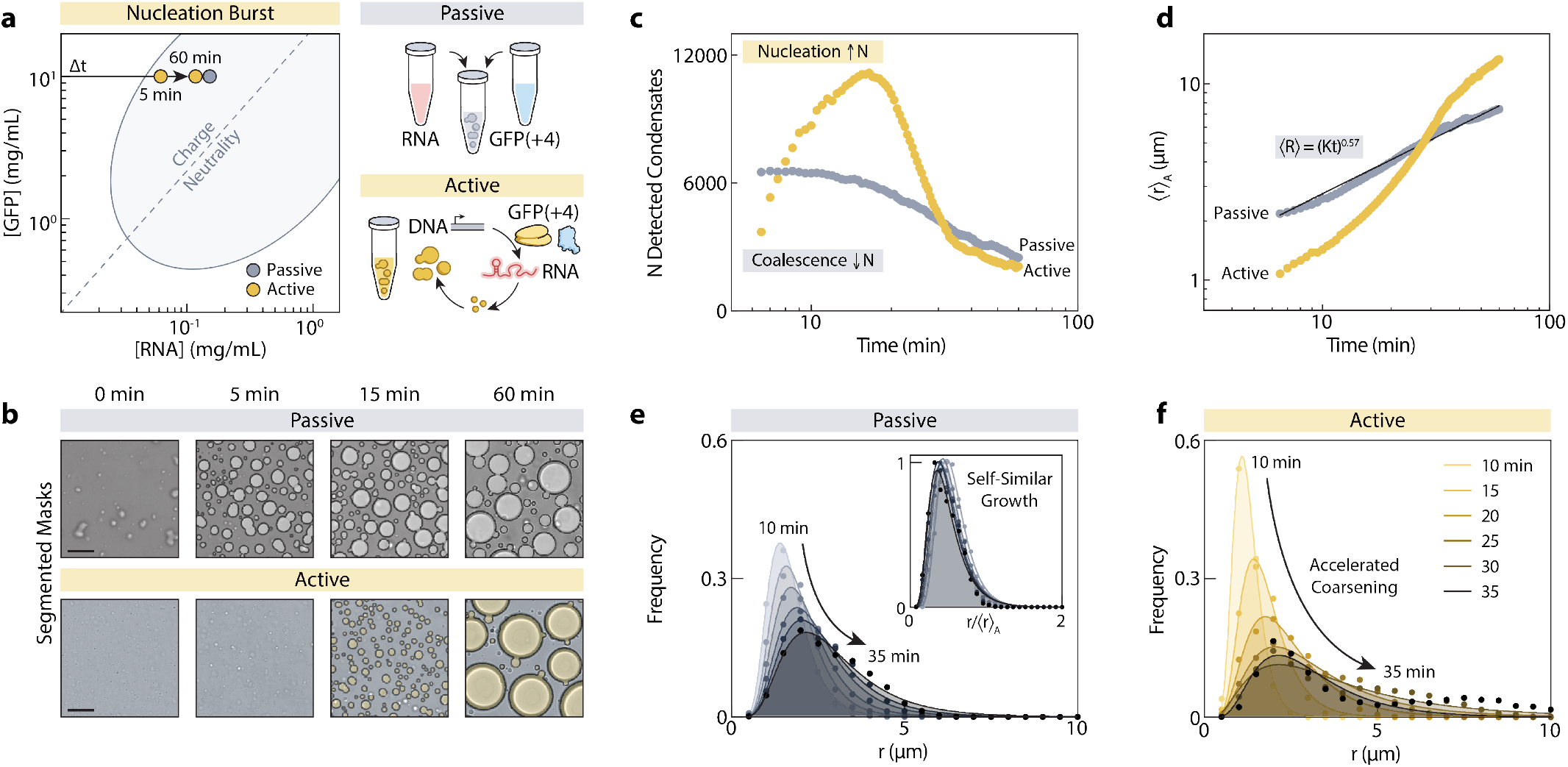
Rapid nucleation burst and accelerated growth kinetics of active condensates. **a.** Phase diagram with the reaction-driven path of the active system and the discrete point of the passive system. Data points correspond to measured RNA concentrations at the indicated reaction times. **b**. Representative microscopy images of passive and active condensates over time with segmented masks overlaid. Scale bars, 20 *µ*m. **c**. Number of detected condensates over time. **d**. Area-weighted mean radius of condensates over time with power-law fit for passive condensates (black line). **e**. Size distributions of passive condensates from 10 to 35 min in 5 min increments. Inset shows distributions rescaled by the mean radius at each time point. **f**. Size distribution of active condensates from 10 min to 35 min in 5 min increments, fit to lognormal distributions. Bin width for panels e and f, 0.5 *µ*m.

Time-course microscopy revealed strikingly different nucleation dynamics between the active and passive cases (Fig. 3b). In the passive system, condensates formed immediately upon mixing and slowly coarsened over time. In the active system, an initial lag phase preceded a rapid nucleation burst, after which condensates quickly coarsened. Image analysis showed that the passive system began with approximately 6,500 detected condensates at the earliest quantifiable time point (6.5 min), with the number of condensates decreasing monotonically over time through fusion and coarsening. In contrast, the active system nucleated more than 11,100 condensates within the first 15.5 minutes before the number of condensates rapidly decreased (Fig. 3c). The area-weighted mean radius, ⟨*r*⟩*A*, in the passive system followed power-law growth consistent with self-similar coarsening, as confirmed by the collapse of rescaled lognormal size distributions onto a single curve (Fig. 3d, e) [36]. This power-law coarsening was consistent both for lower and higher concentrations of purified RNA in the passive system (Supplementary Fig. 8). In contrast, the active system displayed a switch-like response, with size distributions that evolved rapidly between 10 and 35 minutes, unlike the self-similar growth observed in the passive case (Fig. 3f). Interestingly, despite a slightly higher absolute RNA concentration in the passive system, the mean radius of active condensates surpassed the passive counterparts, suggesting that the reaction-driven path of condensate assembly affects the late-stage coarsening behavior. Active condensates formed from RNA transcripts of varying sequence and length all exhibited the same rapid nucleation behavior, suggesting this is a general feature of reaction-driven condensate assembly (Supplementary Fig. 9, 10).

Taken together, our results demonstrate that coupling reaction to phase separation can generate switch-like nucleation dynamics that are qualitatively and quantitatively distinct from passive assembly. This parallels recent work in *C. elegans* demonstrating that transcription-templated assembly, in which nucleation sites are produced dynamically, more accurately captures the rapid nucleolar assembly kinetics observed in early embryos than passive models assuming pre-existing, fixed nucleation sites [13]. More broadly, our findings suggest that the coupling between transcription and phase separation not only regulates condensate size and lifetime, but can also control the timing and rate of nucleation, providing a mechanism for condensates to mount rapid, switch-like transcriptional responses.

### Active condensates regulate transcription through RNA-mediated feedback

Beyond switch-like nucleation, transcriptional condensates have also been proposed to regulate transcription through RNA-mediated feedback, whereby low levels of nascent RNA promote condensate formation and high levels drive dissolution [4]. Although well-supported by cellular observations, direct in vitro reconstitution of both feedback types in a coupled transcription system has remained elusive. Positive feedback was largely inferred from correlations between condensate size and transcriptional output upon addition of a positively charged polyamine, and negative feedback could not be demonstrated in a transcription-coupled system because RNA yields were insufficient to reach the dissolution regime. Our synthetic active system provides a direct approach to test both feedback mechanisms in a coupled transcription system, generating sufficient RNA in situ to traverse the full two-phase region while directly coupling transcription to phase separation.

We first tested whether condensate formation enhances transcriptional output through positive feedback. We performed parallel reactions in the presence and absence of cationic scaffold (2.5 mg mL^-1^) and monitored RNA accumulation over 2 h by SYTO 63 fluorescence (Fig. 4a, b). Although both reactions produced similar levels of RNA initially, the active condensate reaction surpassed the no condensate control after 30 min, yielding approximately 26% more RNA at 2 h. This enhancement was cross-validated by gel densitometry and absorbance at 260 nm after spin column purification, which also confirmed that the additional RNA was of the correct length and recoverable by standard methods (Fig. 4c, Supplementary Fig. 13). We attribute this enhancement to the higher local concentration of T7 RNAP, DNA template, and NTPs within condensates, and it is consistent with prior work showing that macromolecular crowding enhances transcription (Supplementary Fig. 14) [37]. By varying the cationic scaffold and T7 RNAP concentrations, we also demonstrated that RNA yield was most enhanced at a moderate T7 RNAP concentration of 100 nM and scaffold concentrations of 1.5–2.5 mg mL^-1^, corresponding to the broadest accessible region of the phase diagram (Fig. 4d, Supplementary Fig. 14). At lower cationic scaffold concentrations, the two-phase region narrows, so condensates dissolve before transcription enhancement can significantly increase RNA concentration, while at higher concentrations the higher RNA saturation concentration delays condensate formation and the associated transcription enhancement.

**Fig. 4.**
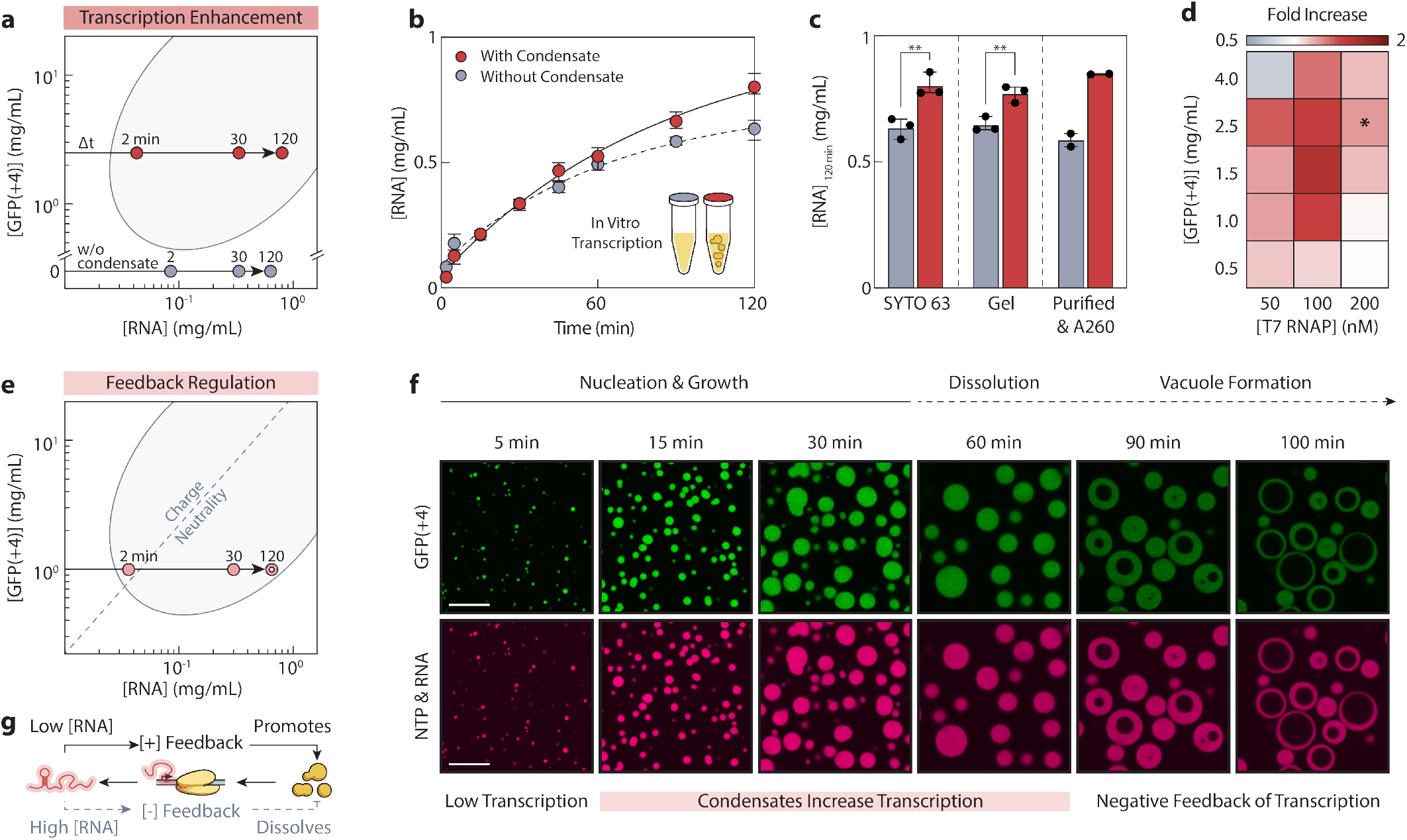
Active condensates regulate transcription through positive and negative RNA-mediated feedback. **a.** Phase diagram and reaction-driven paths for active condensates (200 nM T7 RNAP, 2.5 mg mL^-1^ GFP(+4)) and control reaction without GFP(+4). Data points correspond to measured RNA concentrations at the indicated reaction times. **b**. Total RNA concentration over time in the presence and absence of GFP(+4) condensates as quantified by SYTO 63 fluorescence. Error bars represent the range of 3 independent replicates. **c**. Total RNA concentration at 2 h in the presence and absence of GFP(+4) condensates as quantified by SYTO 63 fluorescence, gel densitometry with SYBR Green II, and absorbance at 260 nm after spin column purification. Data points represent independent replicates, shaded bars represent the mean, and error bars represent the range. P-values were calculated by Welch’s t test (SYTO 63, p = 0.0098; Gel, p = 0.0081). **d**. Fold-change in RNA concentration at 2 h for condensate samples compared to no condensate samples across varying GFP(+4) and T7 RNAP concentrations. Asterisk denotes fold-change for conditions shown in panels a-c. **e**. Phase diagram and reaction-driven path for active condensates at lower GFP(+4) concentration (200 nM T7 RNAP, 1 mg mL^-1^ GFP(+4)). Data points correspond to measured RNA concentrations at the indicated reaction times. **f**. Confocal images showing nucleation, growth, dissolution, and vacuole formation of active condensates corresponding to the reaction-driven path in e., where RNA was labeled with Cy5-UTP during the reaction. Scale bars, 20 *µ*m. **g**. Schematic of positive and negative RNA-mediated feedback, where low RNA concentrations promote condensate formation and excess RNA drives condensate dissolution.

To observe both positive and negative feedback, we then selected conditions that should traverse the full width of the phase diagram, high T7 RNAP concentration (200 nM) and lower cationic scaffold concentration (1 mg mL^-1^) (Fig. 4e). As expected from the phase diagram, condensates formed, grew, and ultimately dissolved as RNA accumulated. Confocal imaging revealed that positive feedback operated during the condensate growth phase, producing a transient enhancement in RNA yield of approximately 26% relative to the no-condensate control at 45 min (Fig. 4f, Supplementary Fig. 7d). Interestingly, while total RNA followed first-order kinetics, RNA concentration in the dilute phase accumulated more slowly and linearly, remaining near the phase boundary throughout active phase separation. This buffering behavior is reminiscent of the proposed role of cellular condensates in stabilizing cytoplasmic concentration against fluctuations [38, 39]. Together, these results support a two-stage feedback model in which moderate RNA concentrations near charge neutrality initially promote transcription by concentrating enzyme and reactants within condensates, before excess RNA pushes the system past the binodal, dissolving condensates and returning transcription to the baseline rate observed in the absence of phase separation (Fig. 4g).

These results suggest that transcription enhancement by condensates may be inherently transient, occurring only in the time window before RNA accumulation drives dissolution. From a condensate engineering perspective, this intrinsic feedback mechanism highlights the importance of understanding reaction-driven paths through the phase diagram to predict when and for how long transcriptional enhancement can occur. Biologically, the duration and magnitude of transcription enhancement could in principle be tuned by modulating scaffold protein and polymerase levels, providing a mechanism to control not just transcriptional enhancement, but the temporal window over which it occurs.

### RNA degradation can renucleate active condensates

Together, our results so far have established that rapid nucleation, RNA-mediated feedback, and transcriptional enhancement can emerge from reaction-phase separation coupling in a minimal active system. We next asked whether this framework could predict even more complex behavior, namely repeated rounds of condensate assembly and dissolution arising from cycles of RNA synthesis and degradation. Based on the passive phase diagram and active reaction-driven paths characterized so far, we hypothesized that after reentrant dissolution, reentry into the two-phase region by RNA degradation could renucleate condensates, giving rise to cyclic condensate behavior resembling a biochemical oscillator.

To test this, we used a high T7 RNAP concentration (200 nM) and a lower cationic scaffold concentration (0.5 mg mL^-1^) to drive RNA accumulation beyond the reentrant arm of the binodal established from the passive phase diagram (Fig. 5a). After 75 min of IVT, we added RNase R, a highly processive 3’ to 5’ exonuclease, to degrade accumulated RNA. Total RNA concentration decayed toward a non-zero plateau of 0.18 mg mL^-1^ with a half-life of approximately 4 min, consistent with a first-order decay model (Fig. 5b). This plateau concentration fell within the two-phase region of the passive phase diagram, meaning the system traversed the two-phase region twice, entering from the low-RNA side during synthesis, exiting through reentrant dissolution as RNA accumulated, and re-entering from the high-RNA side as degradation returned the RNA concentration to the two-phase region (Fig. 5a). The reverse path required exonucleolytic degradation of RNA, as addition of the endonuclease RNase A at varying concentrations or buffer alone did not lead to renucleation of condensates (Supplementary Fig. 15, 16).

**Fig. 5.**
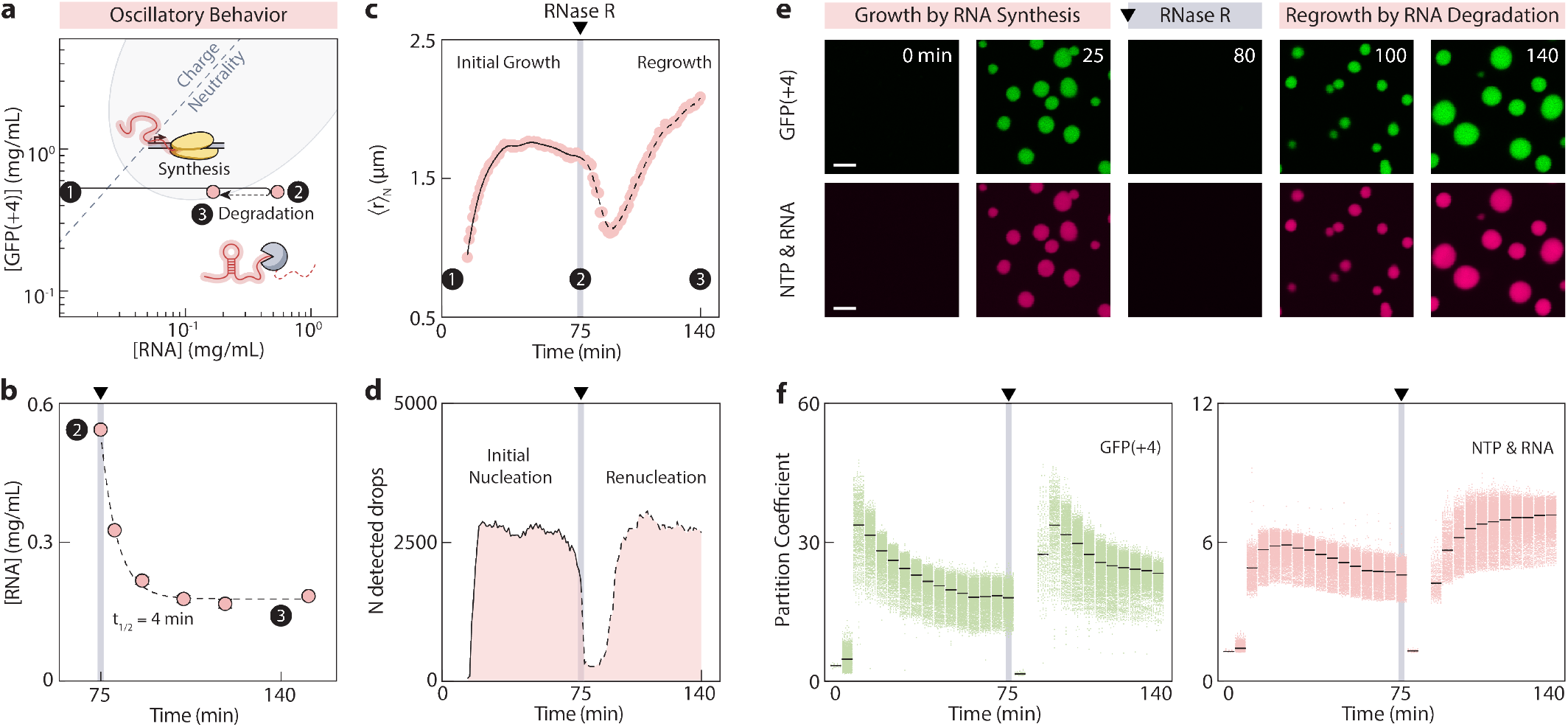
RNA degradation renucleates active condensates. **a.** Phase diagram and the reaction-driven path during RNA synthesis (Step 1 to 2) and degradation (Step 2 to 3) (200 nM T7 RNAP, 0.5 mg mL^-1^ GFP(+4), 20 units RNase R). Data points correspond to measured RNA concentrations at the indicated reaction times. **b**. Total RNA concentration over time quantified by SYTO 63 fluorescence following addition of RNase R at 75 min under the conditions described in a. Concentration was fit to a first-order exponential decay. **c**. Number-weighted mean radius over time during RNA synthesis and degradation. Black line shows LOWESS smoothing with a window of 20 points, with solid and dashed segments indicating the RNA synthesis and degradation phases respectively. **d**. Number of detected condensates over time during RNA synthesis and degradation, with solid and dashed lines indicating the RNA synthesis and degradation phases respectively. **e**. Confocal images showing nucleation, growth, dissolution, and renucleation of active condensates corresponding to the reaction-driven path in a. Scale bars, 5 *µ*m. **f**. Partition coefficients of GFP(+4) (left) and Cy5-UTP (right) in active condensates during the RNA synthesis phase (nucleation and growth) and RNA degradation phase (renucleation and regrowth). Partition coefficient is defined as mean condensate intensity divided by mean dilute phase intensity. Black lines represent the median partition coefficient of all analyzed condensates.

Condensate dynamics during IVT were consistent with phase diagram predictions, with condensates growing during RNA transcription before shrinking at 50 min as excess RNA drove the system toward reentrant dissolution (Fig. 5c). Condensates did not fully dissolve even when the RNA concentration exceeded the passive binodal boundary, suggesting a kinetic barrier to complete dissolution, possibly due to the slow diffusion of RNA out of the dense phase (Supplementary Fig. 17). Addition of RNase R at 75 min dissolved the remaining condensates before renucleating a second generation, with condensate numbers during renucleation closely matching the first generation (∼ 2800 condensates) (Fig. 5d). After RNase R addition, the RNA concentration plateaued within the two-phase region, allowing condensates to equilibrate and grow at near-constant RNA concentration. This led to a mean radius at 140 min that exceeded the maximum mean radius of the first generation. Confocal imaging confirmed successive rounds of condensate nucleation, growth, dissolution, and renucleation (Fig. 5e, f, Supplementary Fig. 18). The RNA and cationic scaffold were enriched to similar extents in both the first and second generation condensates, indicating that condensate compositions follow reproducible paths through the phase diagram.

These results demonstrate that both RNA synthesis and degradation can drive successive rounds of condensate assembly and dissolution, reconstituting the core elements of a biochemical oscillator in a minimal synthetic system. From a biological perspective, scaffold material released upon dissolution could be reused to nucleate a second condensate generation through RNA degradation alone, without requiring additional synthesis. Energetically, this is distinct from the first nucleation event, which is driven by active RNA synthesis and fueled by NTP hydrolysis. Renucleation by degradation is instead a dissipative process that requires no additional energy input and occurs spontaneously as the system returns to charge neutrality.

## Discussion

We demonstrated that RNA flux drives the formation and dissolution of synthetic active condensates along defined paths through the equilibrium phase diagram, recapitulating three dynamic phenomena relevant to cellular transcriptional condensates. First, reaction-phase separation coupling generates a rapid switch-like nucleation burst that is kinetically and quantitatively distinct from passive assembly [13]. The active system nucleated nearly twice as many condensates as the passive system, and these rapid nucleation kinetics shaped late-stage coarsening in ways not captured by equilibrium models. This suggests that the rate and timing of condensate nucleation in cells may be just as functionally important as the final condensate size or composition.

Second, by crossing into and out of the phase boundary through RNA synthesis alone, we reconstituted both positive and negative RNA-mediated feedback regulation in a single coupled experiment, a direct demonstration that was previously inaccessible due to insufficient RNA yields in prior transcription-coupled in vitro systems [4]. Condensate formation enhances transcription by concentrating the reaction machinery, while continued RNA accumulation drives reentrant dissolution, returning transcription to its baseline rate. This feedback is inherently transient and the duration is set by the width of the two-phase region, with important implications for how cells program the temporal window of transcription enhancement.

Third, by exploiting RNase R-driven RNA degradation after dissolution, we demonstrated oscillatory condensate behavior driven by cycles of RNA synthesis and degradation. Degradation-driven renucleation suggests that cells could sustain oscillatory condensate formation without continuously expending energy on new RNA synthesis. These results open several exciting directions at the intersection of non-equilibrium physics, RNA biology, and protocell research. The demonstration that reaction-driven paths through a phase diagram can encode distinct biological functions suggests that cells may possess a richer repertoire of condensate-based regulatory strategies than equilibrium models would predict, programmable simply by tuning the rates of RNA synthesis and degradation. As live-cell imaging approaches reach sufficient spatiotemporal resolution to track condensate nucleation events in real time, and as superresolution methods begin to resolve the molecular composition of condensates during transcriptional bursting, the phase-diagram framework introduced here offers a quantitative basis for interpreting these observations [9, 11, 12, 35].

Perhaps most intriguingly, the cyclic assembly and dissolution of condensates driven by RNA synthesis and degradation raises the possibility of using such systems as platforms for directed evolution under selective pressure [40, 41]. Repeated rounds of condensate formation could selectively concentrate and compartmentalize RNA species that promote their own synthesis or stabilize the condensate, while dissolution and renucleation would provide the competition necessary for selection to act. More broadly, the coupling between chemical reactions and phase separation is emerging as a general organizing principle in cell biology. The minimal synthetic platform we describe provides a tractable experimental foundation for dissecting these phenomena systematically, and for programming life-like temporal organization into protocells that approach the functional complexity of living systems.

## Methods

### Plasmid Construction

Plasmids for GFP(-7) and GFP(0) are available on Addgene (Addgene#199171, Addgene#199166). GFP(-7) contains mutations R90Q and V216A from superfolder GFP [42]. GFP(0) contains mutations E16R, I138R, D207R, N222K, and L246R from GFP(-7). GFP(+4) contains mutations D127R and D86K from GFP(0). DarkGFP(+4) contains mutations of the chromophore-forming residues TYG to GGG. All GFP(+4) mutations were introduced by PCR and confirmed by sequencing. PCR products were purified using a QIAquick PCR Purification Kit (Qiagen) and assembled using NEBuilder HiFi DNA Assembly Master Mix (New England Biolabs). Plasmids were transformed into NiCo21 (DE3) competent cells for protein expression.

### Protein Expression and Purification

The plasmid p6XHis-T7(P266L) was a gift from Anna Pyle (Addgene#174866) [43]. T7 RNAP was expressed and purified from NiCo21 (DE3) cells following adapted methods previously described [44]. Briefly, cells transformed with the plasmid were grown to an optical density (OD600) of 0.7 at 37 °C with shaking at 200 RPM and induced with 0.5 mM isopropyl *ω*-D-1-thiogalactopyranoside (IPTG). After 3 h of expression, cells were harvested by centrifugation and resuspended in lysis buffer (50 mM NaH2PO4, 1 M NaCl, 1 mM DTT pH 8.0). Cells were frozen at -80 °C, thawed, and lysed by sonication at 40% amplitude using a 1/8 inch probe for 10 min with cycles of 2 s on and 4 s off. The lysate was clarified by 2 rounds of centrifugation at 10,000 xg for 30 min each. The protein was purified by immobilized metal affinity chromatography with Ni-NTA resin on an AKTA Start. A 25 mL column was equilibrated with 3 column volumes of lysis buffer prior to sample loading. The column was washed with 10 column volumes of wash buffer (50 mM NaH_2_PO_4_, 1 M NaCl, 35 mM imidazole, pH 8.0) and the T7 RNAP was eluted with 3 column volumes of elution buffer (50 mM NaH_2_PO_4_, 1 M NaCl, 250 mM imidazole, pH 8.0). Fractions were collected during elution, and fractions with absorbance ranging from 100 mAU on the ascending side to 150 mAU on the descending side were pooled. The buffer was then exchanged into 40 mM HEPES, pH 7.4 by spin concentration using spin filters (10 kDa MWCO) with 10 rounds of exchange. The T7 RNAP stock solution was quantified by a Bradford assay and stored at a concentration of 0.42 mg mL^-1^ or 4.1 *µ*M in 50% glycerol, 40 mM HEPES, 100 mM KCl, 1 mM DTT, pH 7.4 at -20 °C.

GFP(+4) was expressed and purified using the method as described for T7 RNAP, but induced with 1 mM IPTG, grown for 16 h post-induction, and purified without DTT. Buffer exchange of purified GFP samples was done at 4 °C by dialysis into 10 mM Tris, pH 7.4 (5 exchanges) followed by 40 mM HEPES, pH 7.4 (1 exchange) with at least 3 h between each exchange. GFP stock solutions were quantified by absorbance at 488 nm with an extinction coefficient of 83,300 M^-1^cm^-1^ and DarkGFP(+4) stock solutions were quantified by Bradford assay (Bio-Rad) and concentrated to 20 mg mL^-1^. Proteins were stored at 4 °C for up to 1 month. All purified proteins were verified by SDS-PAGE.

### Fluorescent Labeling of T7 RNAP

A solution of T7 RNAP (20 *µ*M) in 40 mM HEPES at pH 8 was prepared by exchanging the buffer of the stock solution using a 10 kDa MWCO Ultra Centrifugal Filter. A 20 *µ*L aliquot of Alexa Fluor 594 NHS ester solution in DMSO (2.44 mM) was added to 250 *µ*L of the T7 RNAP solution for a final molar ratio of approximately 10:1 of fluorophore to enzyme. The solution was mixed overnight in a tube rotator at 4 °C and the remaining free dye was separated from the labeled enzyme using 2 Sephadex G-25 HiTrap desalting columns in series on a FPLC (AKTA Start). The eluted fractions were concentrated using 10 kDa MWCO Ultra Centrifugal Filters, quantified by Bradford assay, and stored at a concentration of 0.3 mg mL^-1^ in 40 mM HEPES, 100 mM KCl, 40% glycerol, and 1 mM DTT at -20 °C. As per the manufacturer’s protocol, the absorbance at 590 nm of a 10-fold diluted sample was measured on a NanoDrop and based on the concentration obtained by Bradford assay, the degree of labeling was estimated to be approximately 6.5 or around 10% of total lysines. An extinction coefficient of 73,000 cm^-1^M^-1^ for the Alexa Fluor 594 dye and a molecular weight of 101.4 kDa for T7 RNAP were used.

### Surface Passivation

Glass bottom plates (Cellvis) were passivated following an adapted protocol by etching with 2 M NaOH for 30 min, cleaning with warm 2% Hellmanex for 30 min, and rinsing three times with Milli-Q water [45]. The plate was washed with ethanol and then modified with polyethylene glycol (PEG) using 30 *µ*L of 0.5% mPEG-silane (Laysan Bio mPEG-Silane-5000) in ethanol with 1% acetic acid. This solution was allowed to react with the glass bottom at 70 °C for 20 min. Another 30 *µ*L of mPEG-silane was added and the plate was incubated for an additional 20 min. Wells were rinsed three times with Milli-Q water, dried using compressed air, and incubated at 70 °C for 30 min.

### In Vitro Transcription (IVT)

Linear DNA templates were amplified by PCR with Phusion polymerase, purified with QIAquick spin columns, and quantified by A260 (Thermo Fisher Scientific NanoDrop). Standard transcription reactions were prepared by equilibrating the negatively charged components (DNA template and NTPs) and protein components (cationic GFP and T7 RNAP) separately in 2X HEPES-KCl-MgCl2 buffer before combining to a 1X final buffer concentration. GFP(+4) and T7 RNAP concentrations varied by experiment but were generally in the ranges of 0-10 mg mL^-1^ and 0-200 nM, respectively. The combined reaction was mixed thoroughly by pipetting in a PCR tube and 20 *µ*L was transferred to a 384-well glass-bottom plate for imaging at room temperature (20-22 °C). Unless otherwise specified, the final composition was 76 nM linear DNA template, 40 mM HEPES, 80 mM KCl, 8 mM MgCl2, 6 mM total NTP, pH 7.4. IVT to prepare concentrated RNA for constructing the passive phase diagram was performed with the HiScribe T7 quick high yield RNA synthesis kit (New England Biolabs). Preparative transcription reactions were incubated at 37 °C for 2 h, template DNA was digested with 1 U DNase per 20 *µ*L reaction for 15 min at 37 °C, and the RNA was purified with a Monarch RNA cleanup kit (New England Biolabs).

### RNA Degradation with RNase R

RNA transcription and degradation time-course imaging was performed in a 384-well glass-bottom plate using the standard IVT procedure described above. At the indicated time point, 20 units of RNase R (New England Biolabs) was added to each well and mixed briefly. The plate was resealed with optically transparent film and returned to the microscope for continued imaging. The dilution resulting from RNase R addition was 5% of the total volume.

### Passive Phase Diagram

Stock solutions of 5 mg mL^-1^ of purified RNA and up to 100 mg mL^-1^ of purified GFP(+4) were prepared. Passive phase diagrams of purified RNA and GFP(+4) were prepared by mixing 10 *µ*L of 2X RNA stock in 40 mM HEPES, 80 mM KCl, 8 mM MgCl2, 6 mM ATP, with 10 *µ*L of 2X GFP stock in the same buffer at pH 7.4. The mixtures were then transferred to a 384-well glass-bottom plate and the absorbance at 700 nm was measured with a plate reader (Tecan M200 Pro).

### RNA Quantification and Evaluation

RNA concentration was quantified with SYTO 63 red fluorescent nucleic acid dye (Invitrogen). A large transcription reaction was prepared in a PCR tube and 20 *µ*L was aliquoted into separate tubes for each designated time point. All reactions were incubated at room temperature. Coacervate samples were prepared with DarkGFP(+4) to avoid interfering GFP fluorescence. At each time point, reaction aliquots were quenched with 28 *µ*L of a solution containing HEPES, EDTA, Proteinase K, and urea to a final concentration of 49.4 mM HEPES, 2.5 mM EDTA, 0.4 mg mL^-1^ Proteinase K, and 4.9 mM urea. The quenched reactions were heated to 65 °C for 10 min. The sample was diluted 2 to 16-fold with water to lie within the calibration standard curve (Supplementary Fig. 7a). A sample of 18 *µ*L was transferred to a new PCR tube with 2 *µ*L of 50 *µ*M SYTO 63. The 20 *µ*L sample was transferred to a 384-well glass-bottom plate and the fluorescence intensity was measured by bottom read in a plate reader with excitation at 645 nm, emission at 679 nm, and 100 gain (Tecan M200 Pro). The final dilution factors and the calibration standard curve were used to calculate the RNA concentrations. Transcript length and integrity was analyzed by agarose gel electrophoresis. A 35 mL 2% agarose Tris-borate-EDTA (TBE) gel was prepared with 3.5 *µ*L of 10,000X SYBR Green II (Invitrogen) at a 1X final concentration. A sample of 18 *µ*L from the previously diluted reaction sample was transferred to a new PCR tube and 2 *µ*L of 3 M guanidine hydrochloride was added. RNA loading dye with denaturing agent, formamide, was added to samples and the ssRNA ladder. Samples were heated to 90 °C for 5 min and loaded in the gel. The gel was run at 70 V in 1X TBE buffer (Invitrogen) for 1 h and visualized with a GelDoc gel imaging system (Bio-Rad) with a blue conversion screen and an exposure time of 1 s. Images were exported as 16-bit tif files and analyzed with the Gel Analyzer function in FIJI. As an additional verification of RNA yield, the concentrations of RNA synthesized with and without coacervates were measured by pooling five 20 *µ*L reactions, purifying the RNA using the standard Monarch spin RNA cleanup kit (New England Biolabs) protocol, eluting in 50 *µ*L of water, and quantifying by A260 (Thermo Fisher Scientific NanoDrop).

### Widefield Microscopy

Widefield time-course microscopy images were taken using a Nikon Ti Eclipse microscope. Samples were prepared to a volume of 20 *µ*L in PCR tubes and transferred to a 384-well glass-bottom plate. Images were taken with a 20X Plan-Apochromat NA 0.75 objective lens with excitation at 488 nm for GFP and a Chroma ET - 405/488/561/640 nm quad band filter set. Images were acquired at lower magnification with a sufficiently large field-of-view to sample a large number of condensates for robust statistical analysis. The center of the well was imaged by centering the well using the 4X objective before switching to the 20X objective. The plate was sealed with a transparent film for time-course imaging longer than 2 h.

### Confocal Microscopy

Confocal Laser Scanning Microscopy was performed with a Nikon Ti2 inverted microscope and A1R laser scanning confocal unit with a Plan Apo Lambda 60X/1.40 NA oil immersion objective lens. Fluorescence imaging was performed with sequential excitation at 405 nm, 488 nm, 561 nm, and 640 nm for DAPI, GFP, Alexa Fluor 594-conjugated T7 RNAP, and Cy5-UTP, respectively. Emission filters were 450/50 nm, 525/50 nm, 595/50 nm, and 700/75 nm. Images were acquired using a Galvano scanner with a pinhole set to 1 Airy unit at 640 nm. To minimize fluorescence saturation in the dense phase, mass ratios of 1:100 GFP:DarkGFP, 1:1 labeled:unlabeled T7 RNAP, and 1:50 Cy5-UTP:UTP were used. DNA was pre-incubated with DAPI for 10 min in the dark before adding to the reaction to a final concentration of 3 *µ*M.

### Image Analysis

The image segmentation and quantification pipeline was implemented in Python with the scikit-image library. Condensates from 8-bit image sequences were segmented with Cellpose-SAM (flow threshold = 0.4, cellprob threshold = 0, tile norm blocksize = 200) [46]. Small condensates were segmented with modified parameters of tile norm blocksize=20 and diameter=15. Condensates partially touching the border were removed with clear border. Incorrect masks were removed manually with the object click-select tool in the Cellpose GUI. The properties of labeled objects were quantified with regionprops and plotted in GraphPad Prism. Partition coefficients were estimated from confocal images by segmenting condensates using Otsu thresholding and dividing the mean intensity of each condensate by the mean background intensity. A size pre-filter was set at 20-7000 pixels and condensates partially touching the border were excluded.

## Declarations

### Data availability

Data generated as part of this study are available within the Article and its Supplementary Information and will be available on Figshare at the time of publication.

### Code availability

Python scripts used in this study are available via GitHub at: https://github.com/Obermeyer-Group/Liao_2026.

## Acknowledgments

This work was supported by the National Science Foundation (NSF). J.L. received support from the NSF under award 1848388. Imaging was performed at the Zuckerman Institute’s Cellular Imaging Platform and the Confocal and Specialized Microscopy Shared Resource of the Herbert Irving Comprehensive Cancer Center at Columbia University, funded in part through the NIH/NCI Cancer Center Support Grant P30CA013696. We acknowledge the Precision Biomolecular Characterization Facility at Columbia University for access to the circular dichroism spectrometer supported by NIH Award 1S10OD025102-01.

## Author contribution

J.L., S.Y.A. and A.C.O. conceived the study. J.L. performed the investigations, visualization, and formal analysis. A.C.O. acquired funding. J.L. and A.C.O. wrote the original manuscript draft. J.L., S.Y.A., and A.C.O. reviewed, edited, and approved the final manuscript.

## Competing interests

The authors have filed a provisional patent covering the use of engineered transcriptional condensates to enhance RNA production.

