## Supplementary Information for "Dynamics of synthetic transcriptional condensates emerge from RNA synthesis and degradation"

### Supplementary Methods

**Supplementary Method 1:** Cloning of Additional DNA Templates for IVT
**Supplementary Method 2:** Circular Dichroism (CD)
**Supplementary Method 3:** Fluorescence Characterization
**Supplementary Method 4:** Fluorescence Recovery After Photobleaching (FRAP)

### Supplementary Figures

**Supplementary Fig. 1:** Engineered cationic scaffold proteins and T7 RNAP can be purified by chromatography.
**Supplementary Fig. 2:** Engineered cationic scaffold protein has expected molecular weight and folding.
**Supplementary Fig. 3:** IVT Reaction buffer was optimized to enable transcription-phase separation coupling.
**Supplementary Fig. 4:** Transcription components partition in active condensates.
**Supplementary Fig. 5:** Higher [T7 RNAP] and [GFP(+4)] lead to faster nucleation and growth of condensates.
**Supplementary Fig. 6:** Growth rate of active condensates can be predicted.
**Supplementary Fig. 7:** RNA transcription during active phase separation can be quantified with SYTO 63.
**Supplementary Fig. 8:** Passive condensates with different RNA concentrations all follow power-law coarsening.
**Supplementary Fig. 9:** Active condensates for varying transcript lengths all exhibit rapid nucleation.
**Supplementary Fig. 10:** Active condensates can be assembled with different RNA transcripts.
**Supplementary Fig. 11:** Spectral separation of GFP(+4) and SYTO 63 enables simultaneous quantification.
**Supplementary Fig. 12:** The dilute phase composition remains near the phase boundary during phase separation.
**Supplementary Fig. 13:** RNA from active condensates was quantified and evaluated by gel densitometry.
**Supplementary Fig. 14:** Active condensates concentrate T7 RNAP and enhance transcription.
**Supplementary Fig. 15:** RNase R can renucleate condensates.
**Supplementary Fig. 16:** Buffer and RNase A do not renucleate active condensates after reentrant dissolution.
**Supplementary Fig. 17:** GFP(+4) is more mobile than RNA within active condensates.
**Supplementary Fig. 18:** Dense phase compositions oscillate during transcription and degradation.

### Supplementary Tables

**Supplementary Table. 1:** Physical Parameters of Engineered Cationic Scaffold and RNA
**Supplementary Table. 2:** Active Condensate Growth and Dissolution Reaction Conditions
**Supplementary Table. 3:** Active and Passive Condensate Reaction Conditions
**Supplementary Table. 4:** Transcription and Degradation Reaction Conditions
**Supplementary Table. 5:** Generalization of Active Condensates to Different RNA
**Supplementary Table. 6:** Primers for Generating Linear DNA Templates for IVT
**Supplementary Table. 7:** Key Resources Table

### Supplementary Videos

**Supplementary Video. 1:** Active condensate growth increases with T7 RNAP concentration
**Supplementary Video. 2:** Active condensate growth increases with cationic scaffold concentration
**Supplementary Video. 3:** Active condensates form vacuoles during reentrant phase transitions
**Supplementary Video. 4:** Active condensates show rapid nucleation burst and accelerated growth kinetics
**Supplementary Video. 5:** Active condensates exhibit positive and negative RNA-mediated feedback
**Supplementary Video. 6:** Active condensates renucleate by RNA degradation
**Supplementary Video. 7:** FRAP of GFP(+4) and Cy5-UTP in active condensates

### Cloning of Additional DNA Templates for IVT

DNA templates encoding different RNA transcripts were cloned into a pET24a(+) vector backbone. $ELP_{S96}$ encoding 96 repeats of an elastin-like polypeptide with serine at the guest residue position in a pET25b(+) vector was obtained from Addgene (Plasmid #68393). Shorter ELP variants, $ELP_{S6}$ and $ELP_{S18}$, were PCR-amplified from this template and inserted into pET24a(+) by HiFi assembly. The gene for the disordered region of the nucleosome linker histone H5 (derived from PDB: 7XVM) was synthesized in pET24a(+) by TWIST Bioscience. Both the shorter variant $H5_{2/3}$ and the tandem repeat construct H5-H5 were generated by amplifying H5 and assembling into pET24a(+) using HiFi assembly. Four repeats of a Pepper aptamer (FR Biotechnology) were inserted downstream of the gene of interest before the T7 terminator by PCR amplification and HiFi assembly, generating GFP(0)-4Pepper and H5-4Pepper.

### Circular Dichroism (CD)

CD spectra were obtained at room temperature using a 1 mm quartz cuvette (Hellma) and a Chirascan V100 CD spectrometer. Protein samples in 40 mM HEPES, pH 7.4 were diluted 80-fold to 0.25 mg/mL with MilliQ water prior to measurement. Spectra were collected from 300 to 200 nm in 1 nm steps with an integration time of 0.2 s.

### Fluorescence Characterization

Fluorescence excitation and emission spectra for GFP(+4) and SYTO 63 were measured at room temperature using a plate reader (Tecan M200 Pro). For GFP(+4), protein was prepared at 0.5 mg $mL^{-1}$ in 40 mM HEPES, pH 7.4. Spectra were collected with excitation at 420 nm (for emission scans) and emission at 580 nm (for excitation scans), with 4 nm step sizes at a gain of 80 by bottom read. Emission spectra of sfGFP and DarkGFP(+4) were also collected under identical conditions. For SYTO 63, 2 µL of a 50 µM stock solution was added to 18 µL of purified RNA (0.5 mg $mL^{-1}$) to a final SYTO 63 concentration of 5 µM. Spectra were collected with excitation at 600 nm (for emission scans) and emission at 750 nm (for excitation scans), with 4 nm step sizes at a gain of 100 by bottom read.

### Fluorescence Recovery After Photobleaching (FRAP)

Confocal imaging was performed with a Nikon Ti2 inverted microscope and AX R laser scanning confocal unit with a 60X/1.49 NA Apochromat oil immersion objective lens. Fluorescence imaging was performed with sequential excitation at 488 nm (524±18 nm emission) for GFP and 640 nm (699.5±37.5 nm emission) for Cy5-UTP. FRAP was performed on condensates in a 256x256 field of view with a pixel size of 71.9 nm by selecting 1 $\mu$m diameter regions of interest (ROIs) for reference and bleaching in the NIS Elements software. Nine pre-bleach frames were acquired before samples were sequentially bleached with 2% 488 nm laser and 8% 561 nm laser with a dwell time of 2 $\mu$s. The post-bleach recovery acquisition rate was 1 frame per 3 seconds. FRAP recovery profiles were quantified in FIJI, double-normalized using background and unbleached reference regions, scaled to 0–1, and fitted to a single or bi-exponential.

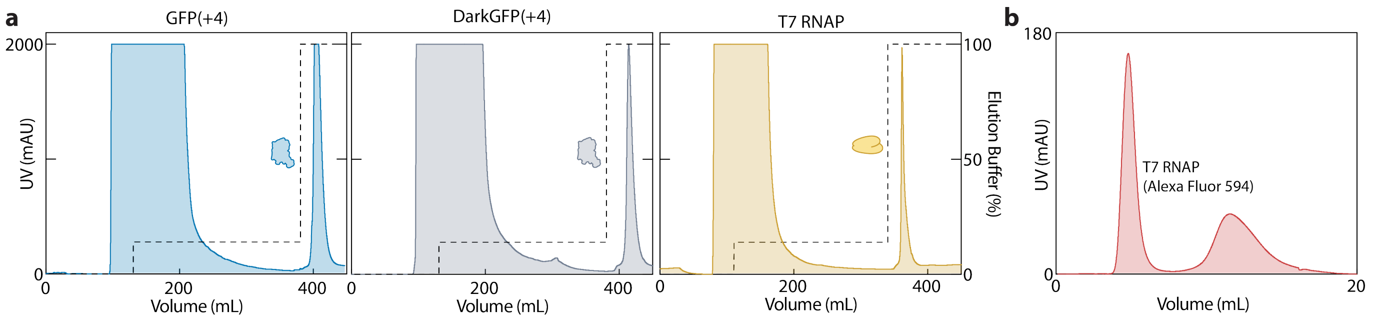


**Supplementary Fig. 1 Engineered cationic scaffold proteins and T7 RNAP can be purified by chromatography. a.** Nickel affinity chromatograms of GFP(+4), DarkGFP(+4), and T7 RNAP showing flow-through and elution peaks. Dashed line indicates the percentage of elution buffer, with steps at 14% (wash) and 100% (elution). **b.** Desalting chromatogram of T7 RNAP labelled with Alexa Fluor 594.

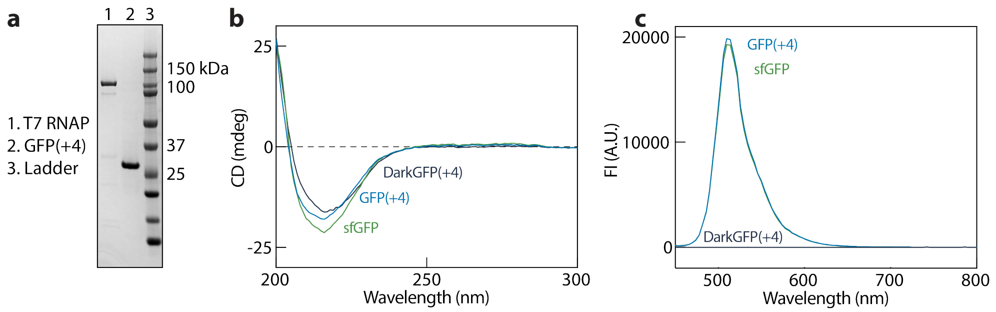


**Supplementary Fig. 2 Engineered cationic scaffold protein has expected molecular weight and folding. a.** SDS-PAGE gel of T7 RNAP and GFP(+4). Expected molecular weight of T7 RNAP and GFP(+4) are 101.4 and 28.01 kDa, respectively. **b.** Circular dichroism spectra of sfGFP, GFP(+4), and DarkGFP(+4). **c.** Fluorescence emission spectra of sfGFP, GFP(+4), and DarkGFP(+4) with excitation at 420 nm.

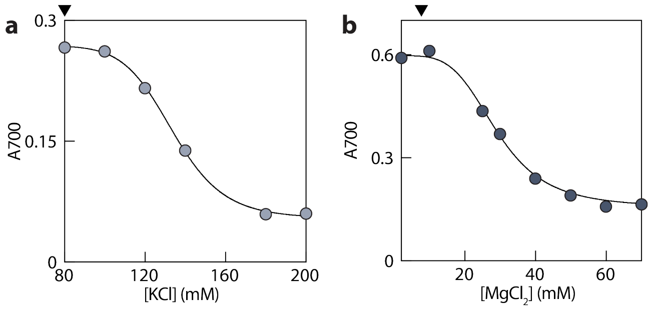


**Supplementary Fig. 3 IVT Reaction buffer was optimized to enable transcription-phase separation coupling. a.** Turbidity of passive condensates at increasing concentrations of KCl. **b.** Turbidity of passive condensates at increasing concentrations of $MgCl_2$. Conditions for a-b were 1 mg $mL^{-1}$ GFP(+4), 0.5 mg $mL^{-1}$ RNA, 40 mM HEPES, pH 7.4. GFP(+4) and RNA were pre-equilibrated separately in each buffer before mixing. Black lines show sigmoidal four-parameter logistic fit. Black triangles indicate the standard concentrations of 80 mM KCl and 8 mM $MgCl_2$ chosen for the IVT reaction buffer.

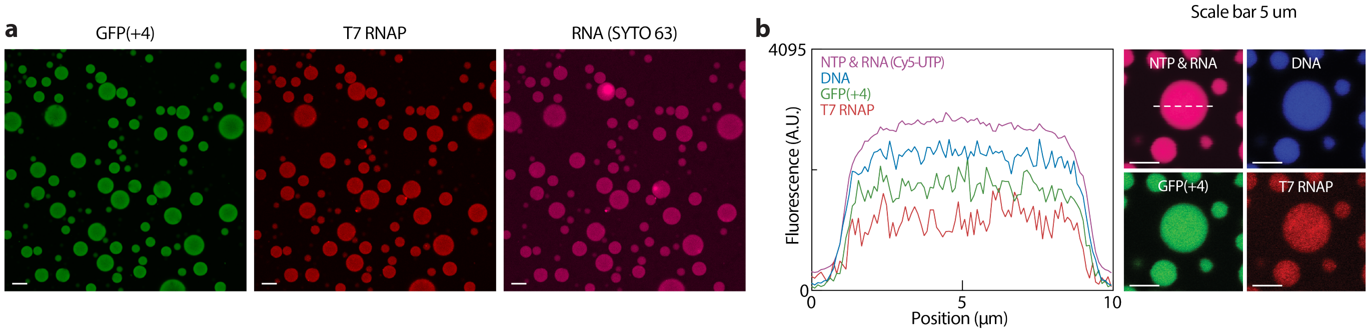


**Supplementary Fig. 4 Transcription components partition in active condensates. a.** Confocal images showing the co-partitioning of GFP(+4), T7 RNAP (Alexa Fluor 594), and RNA (SYTO 63) within an active condensate after 60 min of IVT. Total DarkGFP/GFP and T7 RNAP concentrations were 2.5 mg $mL^{-1}$ and 50 nM, respectively. Scale bars, 20 $\mu$m. **b.** Confocal images corresponding to Fig. 1e showing the co-partitioning of T7 RNAP (Alexa Fluor 594), linear template DNA (DAPI), NTP and RNA (Cy5-UTP), and GFP(+4) within an active condensate after 60 min of IVT. Total DarkGFP/GFP and T7 RNAP concentrations were 1 mg $mL^{-1}$ and 100 nM, respectively. Scale bars, 5 $\mu$m. Reaction condition for a-b was 76 nM linear DNA template, 40 mM HEPES (pH 7.4), 80 mM KCl, 8 mM $MgCl_2$, and 6 mM total NTP. Mass ratios of 1:100 GFP:DarkGFP, 1:1 labeled:unlabeled T7 RNAP, and 1:50 Cy5-UTP:UTP were used.

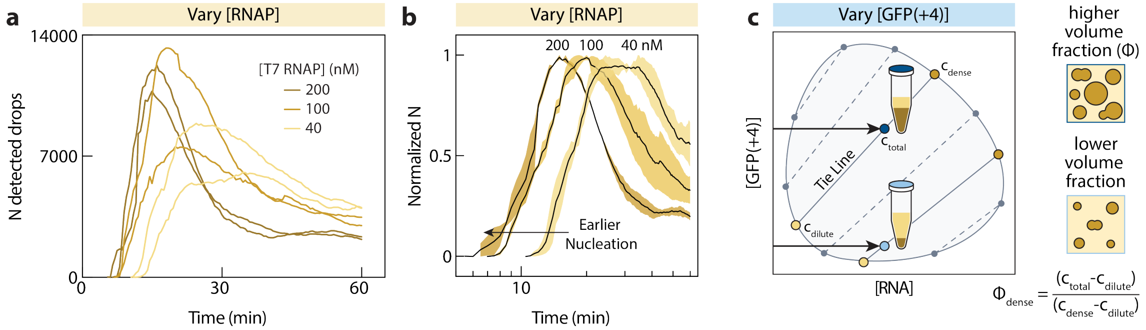


**Supplementary Fig. 5 Higher [T7 RNAP] and [GFP(+4)] lead to faster nucleation and growth of condensates. a.** Total number of detected condensates for varying T7 RNAP concentrations. **b.** Normalized number of detected condensates for varying T7 RNAP concentrations. Shaded regions indicate range of independent duplicates. Black line denotes the mean. Reaction conditions are the same as Fig. 2a-c. **c.** Schematic of hypothesized phase diagram with tie lines and reaction-driven paths at higher and lower [GFP(+4)].

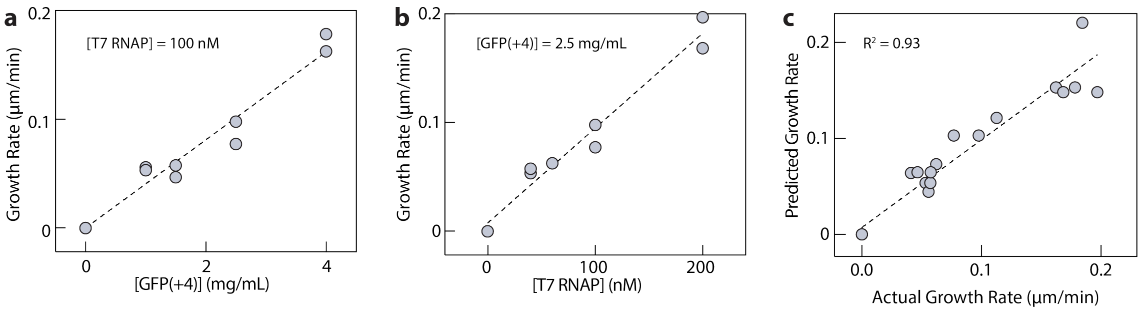


**Supplementary Fig. 6 Growth rate of active condensates can be predicted a** Growth rate between 20-30 min increases with [GFP(+4)] at constant [T7 RNAP] of 100 mM. **b** Growth rate between 20-30 min increases with [T7 RNAP] at constant [GFP(+4)] of 2.5 mg mL$^{-1}$. **c** Actual and predicted growth rate of active condensates between 20-30 min for varying T7 RNAP and GFP(+4) concentrations, corresponding to Fig. 2f. The increase in $\langle r \rangle_N$ between 20 and 30 min from Fig. 2f was estimated by linear regression and then fit to a multiplicative saturation model where,

$$\text{Growth Rate, } \frac{d\langle r \rangle_N}{dt} = V_{\max} \left( \frac{[\text{T7RNAP}]}{K_{T7RNAP} + [\text{T7RNAP}]} \right) \left( \frac{[\text{GFP(+4)}]}{K_{GFP(+4)} + [\text{GFP(+4)}]} \right) \tag{1}$$

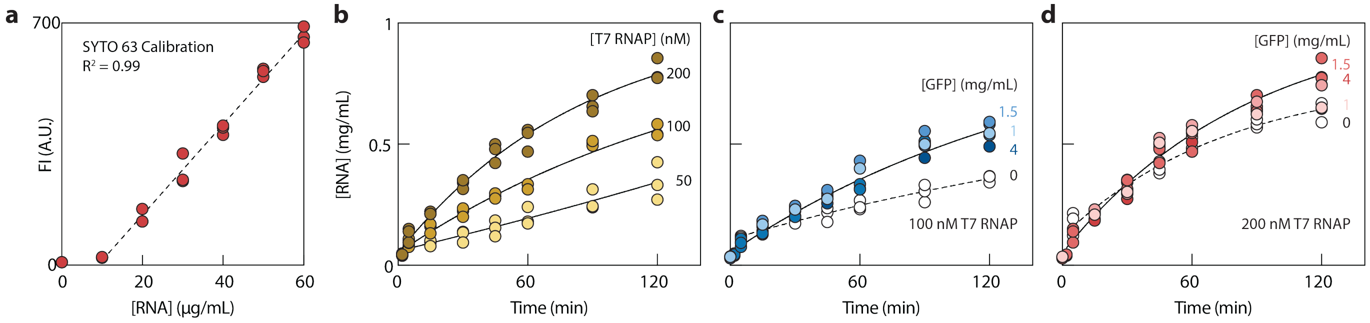


**Supplementary Fig. 7 RNA transcription during active phase separation can be quantified with SYTO 63. a.** Linear calibration curve for purified RNA quantified by SYTO 63 fluorescence. Total RNA concentration over time for conditions corresponding to **b.** Fig. 2a-b, **c.** Fig. 2d-e, and **d.** Fig. 2g-i. As a guide to the eye, data were approximated by a first-order kinetic model with effective transcription rate constant $k_t$, degradation rate constant $k_d$, and baseline b.

$$[RNA] = \frac{k_t}{k_d} \left( 1 - e^{-k_d t} \right) + b \tag{2}$$

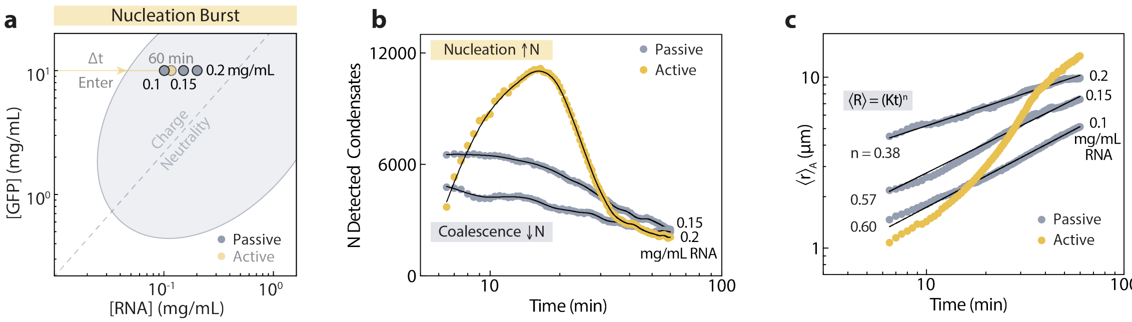


**Supplementary Fig. 8 Passive condensates with different RNA concentrations all follow power-law coarsening. a.** Phase diagram and the discrete RNA concentrations corresponding to three passive reference conditions. **b.** Number of detected condensates over time for active system and passive system with 0.15 and 0.2 mg mL$^{-1}$ purified RNA. Black line shows LOWESS smoothing with 20 points smoothing window. **c.** Area-weighted mean radius of passive condensates over time with power-law fit (black line) at varying concentrations of purified RNA compared to active condensates (yellow).

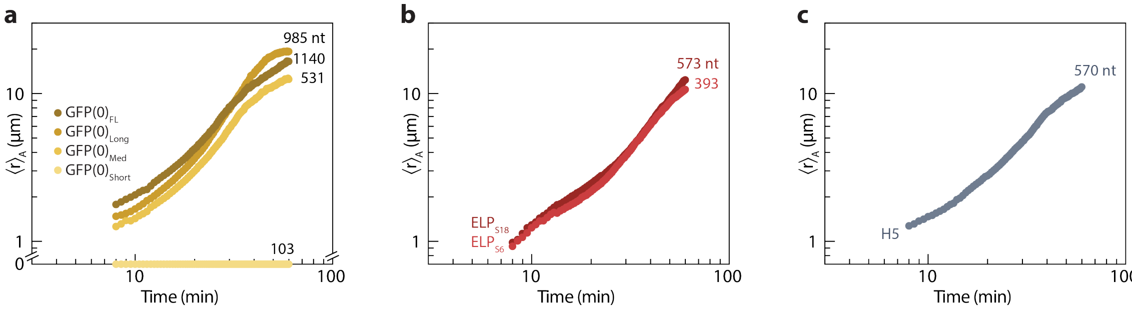


**Supplementary Fig. 9 Active condensates for varying transcript lengths all exhibit rapid nucleation.** Area-weighted mean radius of active condensates formed with RNA transcripts of varying length, transcribed from equimolar DNA templates encoding variants of **a.** GFP(0), **b.** elastin-like polypeptide with a serine guest residue, and **c.** histone H5, respectively as described in the Supplementary Methods.

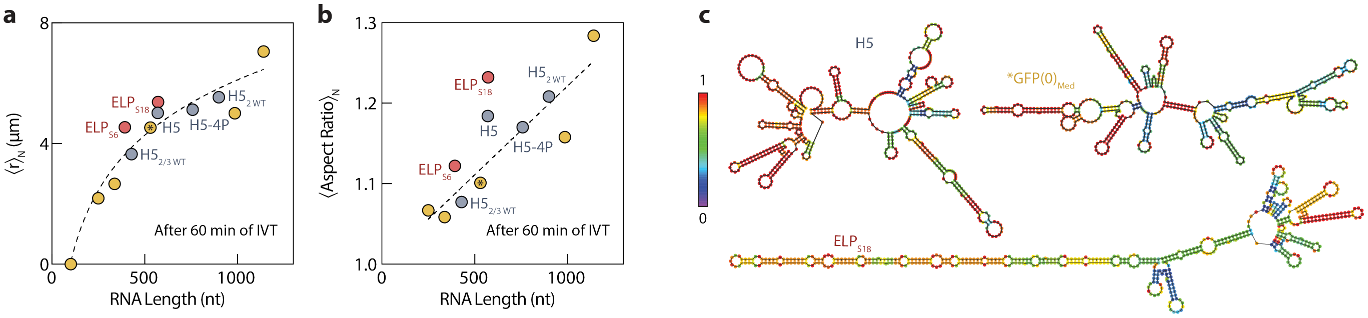


**Supplementary Fig. 10 Active condensates can be assembled with different RNA transcripts. a.** Number-weighted mean radius of active condensates after 60 min of IVT, for transcripts encoding GFP(0) (yellow), elastin-like polypeptide with a serine guest residue (red), and the disordered region of histone H5 (gray). Mean radius increases with RNA length. Semilog line shown as guide to the eye. **b.** Number-weighted mean aspect ratio of active condensates increases with RNA length. Asterisk indicates the RNA characterized in the main text. **c.** Minimum free energy secondary structures predicted by RNAfold, colored by base-pair probability.

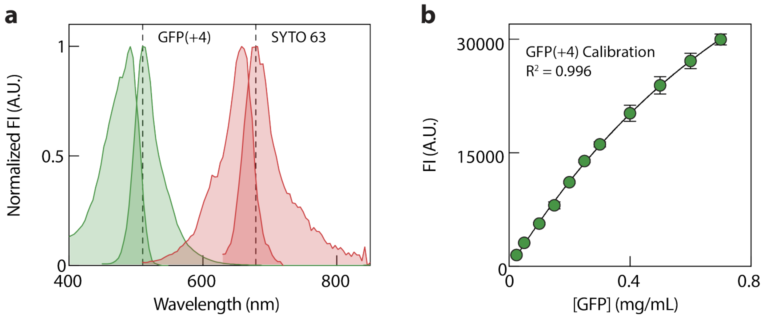


**Supplementary Fig. 11 Spectral separation of GFP(+4) and SYTO 63 enables simultaneous quantification. a.** Normalized fluorescence excitation and emission spectra for GFP(+4) and SYTO 63 as described in the Supplementary Methods. Dashed lines denote wavelength near max emission used for quantification. **b.** Fluorescence calibration curve for GFP(+4) (470/510 nm, gain 70, bottom read). Black line is an exponential plateau fit.

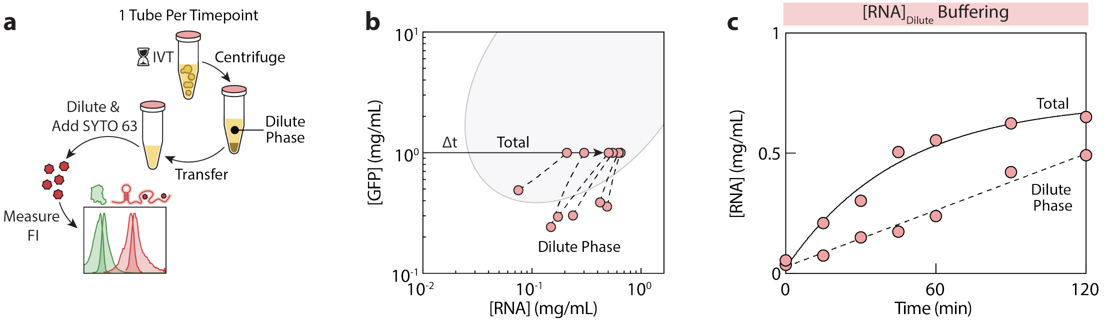


**Supplementary Fig. 12 The dilute phase composition remains near the phase boundary during phase separation. a.** Schematic of dilute phase quantification during IVT with active condensates. Independent reactions for each time point were centrifuged at 4000 xg and the supernatant was removed, diluted, and quantified by fluorescence. **b.** Total and dilute phase compositions during IVT with active condensates, overlaid on the passive phase boundary. Dashed lines approximate partial tie lines. Circles are time points sampled over 120 min. Reaction conditions correspond to Fig. 4e. **c.** Total and dilute phase RNA concentration over time. As a guide to the eye, the total concentration and dilute phase were fit to a first-order kinetic model and linear regression, respectively.

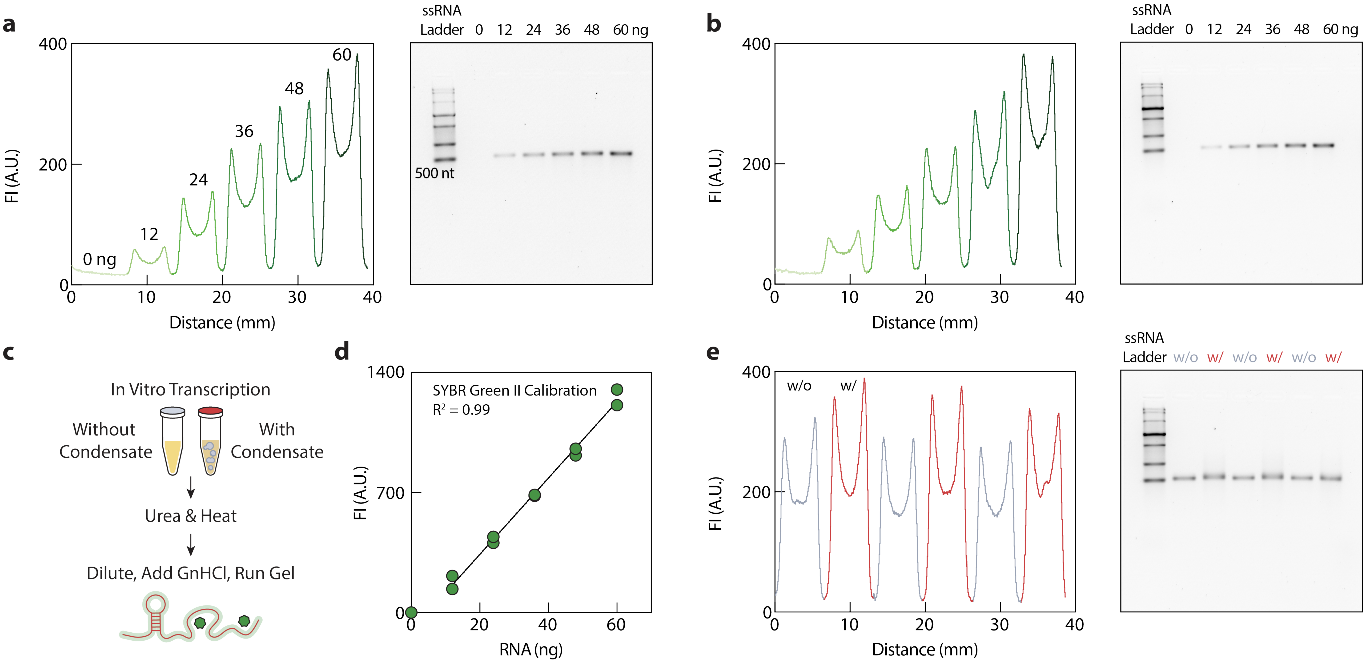


**Supplementary Fig. 13 RNA from active condensates was quantified and evaluated by gel densitometry. a.** Gel densitometry analysis of agarose gel with varying amounts of purified RNA for calibration. **b.** Replicate of gel densitometry calibration. **c.** Schematic of quantification method with urea and heat dissolution of condensates and gel densitometry by SYBR Green II fluorescence. **d.** Linear calibration curve of RNA concentration from gel densitometry. **e.** Quantification and evaluation of RNA from samples without and with active condensates corresponding to the conditions of Fig. 4c.

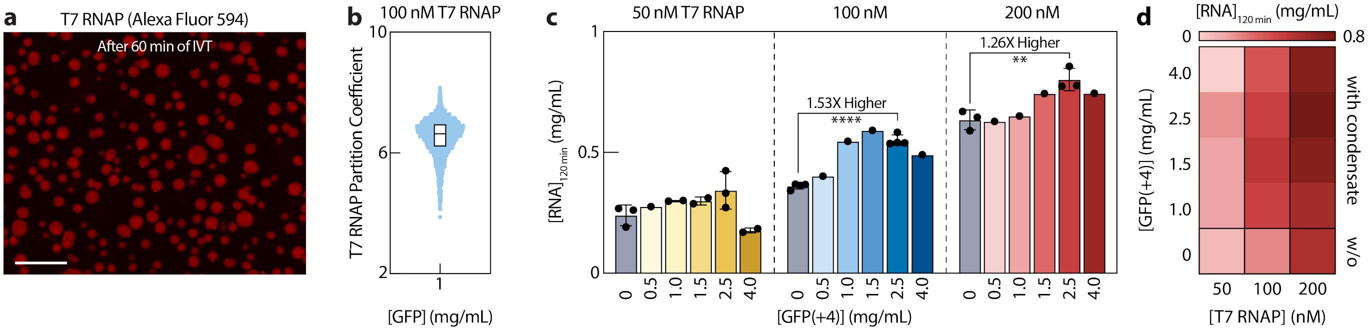


**Supplementary Fig. 14 Active condensates concentrate T7 RNAP and enhance transcription. a.** Confocal image showing the co-partitioning of T7 RNAP (Alexa Fluor 594) within active condensates after 60 min of IVT. Scale bar, 20 $\mu$m. **b.** Partition coefficient of T7 RNAP (Alexa Fluor 594) within active condensates after 60 min of IVT. Data points represent the mean partition coefficient of individual condensates. Box plot represents the median and interquartile range. **c.** Total RNA concentration after 2 h of IVT for varying GFP(+4) and T7 RNAP concentrations. Bar graph with individual data points. Where appropriate, error bars represent the range. P-values were calculated by Welch's t test (**** is $p < 0.0001$ and ** is $p = 0.0098$). **d.** Heat map of the same data shown in c.

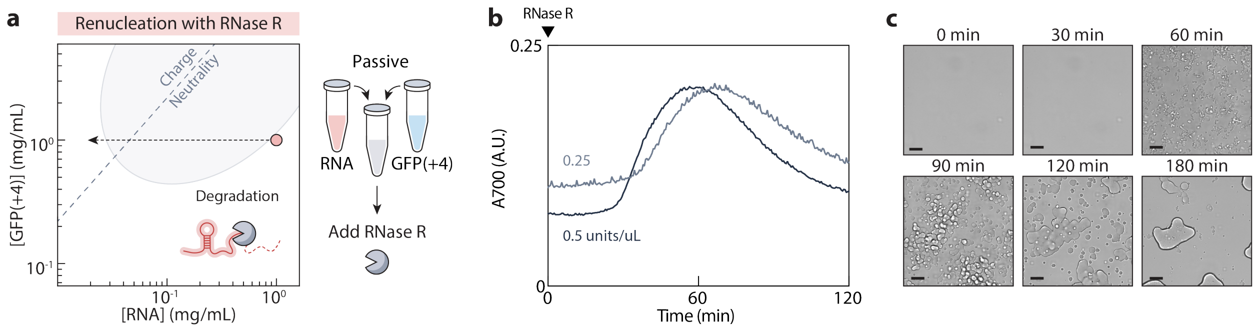


**Supplementary Fig. 15 RNase R can renucleate condensates. a.** Schematic of phase diagram and the RNase-driven path during RNA degradation (1 mg mL$^{-1}$ GFP(+4), 1 mg mL$^{-1}$ purified RNA). Buffer condition was 40 mM HEPES (pH 7.4), 80 mM KCl, 8 mM $MgCl_2$, and 6 mM total NTP. RNase R was added immediately after mixing GFP(+4) and RNA. **b.** Turbidity of RNA degradation-driven renucleation of condensates for 0.25 and 0.5 units/$\mu$L RNase R. **c.** Microscopy images over time during renucleation and dissolution of condensates during RNA degradation. Condition was 1.5 mg mL$^{-1}$ GFP(+4), 1.6 mg mL$^{-1}$ purified RNA, and 0.5 units/$\mu$L RNase R in 40 mM HEPES (pH 7.4), 80 mM KCl, 8 mM $MgCl_2$, and 6 mM total NTP. Scale bars, 20 $\mu$m.

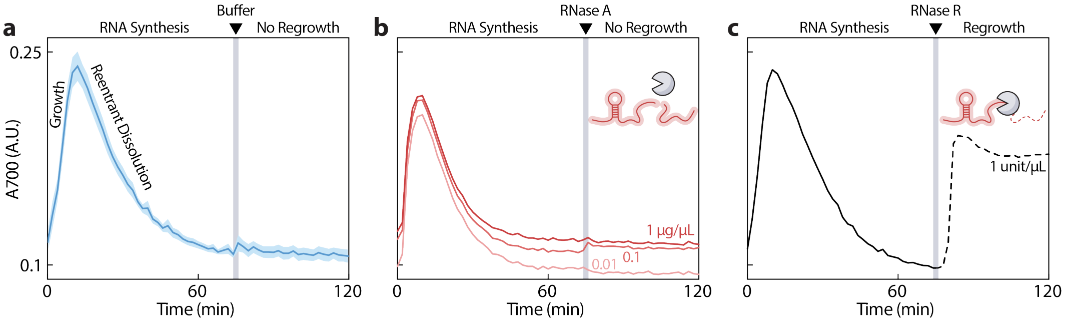


**Supplementary Fig. 16 Buffer and RNase A do not renucleate active condensates after reentrant dissolution. a.** Turbidity of active condensates showing formation by transcription, reentrant dissolution, and no renucleation upon addition of 40 mM HEPES buffer. **b.** Turbidity of active condensates showing formation by transcription, reentrant dissolution, and no renucleation at varying RNase A concentrations. **c.** Turbidity of active condensates showing formation by transcription, reentrant dissolution, and renucleation upon addition of 20 U RNase R. Reaction conditions for a–c correspond to Fig. 5, except for the buffer or RNase addition. Gray lines indicate time of addition.

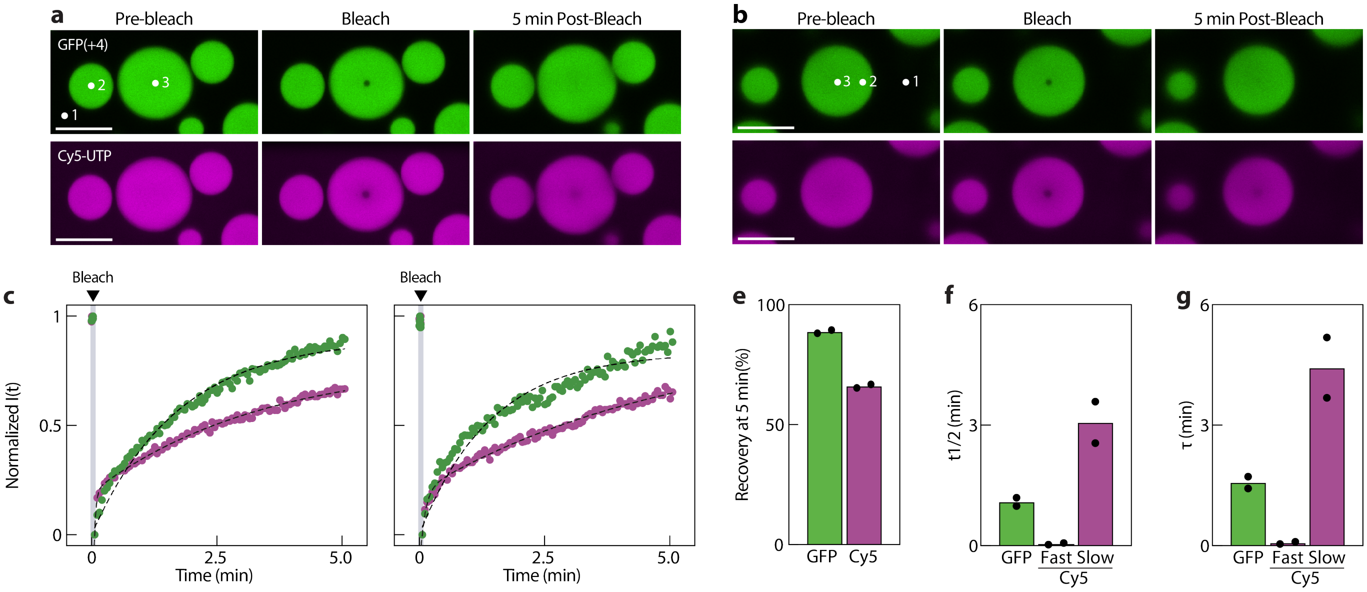


**Supplementary Fig. 17 GFP(+4) is more mobile than RNA within active condensates. a-b.** Representative FRAP confocal images showing background (1), unbleached reference (2), and bleached (3) regions of interest. Scale bars, 10 $\mu$m. Images are from independent active condensates after 1.2-1.5 h of IVT. Reaction conditions were 100 nM T7 RNAP, 2.5 mg mL$^{-1}$ GFP(+4), with 1:100 GFP(+4):DarkGFP(+4), 1:1 labeled:unlabeled T7 RNAP, and 1:50 Cy5-UTP:UTP. **c-d.** FRAP recovery curves for GFP(+4) (green) and Cy5-UTP-labeled NTP and RNA (purple). Gray line indicates the bleach point. Dashed lines are exponential fits, where GFP(+4) was fit to a single exponential and Cy5-UTP to a bi-exponential, with the fast and slow components corresponding to free UTP and RNA, respectively. **e-g.** Recovery at 5 min, half-time, and time constant from single and bi-exponential fits for GFP(+4) (green) and RNA (Cy5-UTP, purple), respectively.

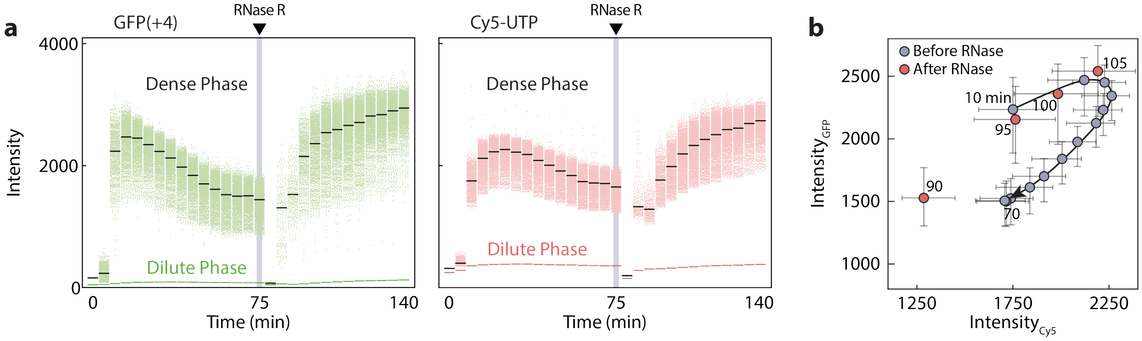


**Supplementary Fig. 18 Dense phase compositions oscillate during transcription and degradation. a.** Fluorescence intensities of GFP(+4) (left) and Cy5-UTP (right) in the dilute and dense phases quantified by confocal microscopy. Data points represent the mean intensity of individual condensates. Black lines indicate the median of individual condensate mean intensities. Conditions correspond to Fig. 5. RNase R (20 units) was added at 75 min. Intensity values are on a 12-bit scale (0–4095). **b.** Fluorescence intensities of GFP(+4) and Cy5-UTP in the dense phase over time before and after RNase R addition. Markers and error bars represent the median and interquartile range.

**Table 1** Physical Parameters of Engineered Cationic Scaffold and RNA

| Parameter | GFP(+4) | RNA |
|---|---|---|
| $M$ (kDa)[1] | 28.01 | 171.3 |
| $N$ (a.a. or nt.) | 248 | 531 |
| $Q$ (at pH 7.4) | +3.97 | -531 |

[1]M of GFP includes methionine. M of RNA is for 5' triphosphate.

**Table 2** Active Condensate Growth and Dissolution Reaction Conditions

| Figure | Replicate | Date | [T7 RNAP] (nM) | [GFP(+4)] (mg mL$^{-1}$, $\mu$M) |
|---|---|---|---|---|
| Fig. 2b | 1 | 2025.10.14 (Run 1) | 40 | 2.5, 89 |
| | 2 | 2025.10.26 (Run 2) | | |
| | 1 | 2025.10.15 (Run 1) | 100 | |
| | 2 | 2025.12.28 (Run 1) | | |
| | 1 | 2025.10.26 (Run 1) | 200 | |
| | 2 | 2025.12.27 (Run 1) | | |
| Fig. 2d | 1 | 2025.10.15 (Run 2) | 100 | 1.0, 36 |
| | 2 | 2026.04.26 (Run 1) | | |
| | 1 | 2025.11.02 (Run 2) | | 1.5, 54 |
| | 2 | 2025.12.30 (Run 1) | | |
| | 1 | 2025.10.15 (Run 1)[1] | | 2.5, 89 |
| | 2 | 2025.12.28 (Run 1)[1] | | |
| | 1 | 2025.11.01 (Run 1) | | 4.0, 143 |
| | 2 | 2025.12.28 (Run 2) | | |
| Fig. 2 f (Additional Data) | 1 | 2025.10.14 (Run 2) | 60 | 2.5, 89 |
| | 1 | 2025.10.20 (Run 1) | 200 | 1.0, 36 |
| | 1 | 2025.11.01 (Run 2) | 200 | 2.0, 71 |
| | 1 | 2025.12.27 (Run 1) | 200 | 2.5, 89 |
| | 1 | 2025.12.29 (Run 1) | 200 | 4, 143 |
| Fig. 2h | 1 | 2025.11.03 (Run 1) | 200 | 1.5, 54 |
| Fig. 2i | 1 | 2025.10.20 (Run 1) | 200 | 1.0, 36 |
| | 2 | 2025.11.06 (Run 1) | | |
| | 1 | 2025.10.27 (Run 1) | | 1.5, 54 |
| | 2 | 2025.11.03 (Run 1)[2] | | |
| | 1 | 2025.11.02 (Run 1) | | 2.0, 71 |
| | 1 | 2025.11.22 (Run 1) | | 2.5, 89 |

Reaction buffer was 76 nM linear DNA, 40 mM HEPES, 80 mM KCl, 8 mM $MgCl_2$, 6 mM total NTP, pH 7.4

[1]Same data as Fig. 2b

[2]Same data as Fig. 2h

**Table 3** Active and Passive Condensate Reaction Conditions

| Figure | Date | [T7 RNAP] (nM) | [Purified RNA] (mg mL$^{-1}$, $\mu$M) | [GFP(+4)] (mg mL$^{-1}$, $\mu$M) |
|---|---|---|---|---|
| Fig. 4 (Passive) | 2026.01.24 (Run 2) | 0 | 0.10, 0.58 | 10, 357 |
| | 2026.01.26 (Run 1) | | 0.15, 0.88 | |
| | 2026.01.24 (Run 3) | | 0.20, 1.17 | |
| Fig. 4 (Active) | 2026.01.15 (Run 3) | 50 | 0 | 10, 357 |

Buffer for passive and active condensates was 76 nM linear DNA, 40 mM HEPES, 80 mM KCl, 8 mM $MgCl_2$, 6 mM total NTP, pH 7.4.

**Table 4** Transcription and Degradation Reaction Conditions

| Figure | Date | [T7 RNAP] (nM) | [RNase R] (Units) | [GFP(+4)] (mg mL$^{-1}$, $\mu$M) |
|---|---|---|---|---|
| Fig. 5c-d | 2026.03.31 (Run 2) | 200 | 20 | 0.5, 18 |
| Fig. 5e-f | 2026.04.11 (Run 2) | 200 | 20 | 0.5, 18 |

Reaction buffer was 76 nM linear DNA, 40 mM HEPES, 80 mM KCl, 8 mM $MgCl_2$, 6 mM total NTP, pH 7.4. RNase R was added at approximately 75 min.

**Table 5** Generalization of Active Condensates to Different RNA

| Figure | Date | RNA Template | RNA Length (nt) |
|---|---|---|---|
| Supplementary Fig. 9 | 2026.01.14 (Run 3) | GFP(0) | 103 |
| | 2026.02.21 (Run 1) | | 250 |
| | 2026.02.22 (Run 1) | | 338 |
| | 2026.01.15 (Run 3) | | 531 |
| | 2026.01.15 (Run 4) | | 985 |
| | 2026.02.01 (Run 2) | GFP(0)-4Pepper | 1140 |
| | 2026.01.31 (Run 2) | $ELP_{S6}$ | 393 |
| | 2026.01.31 (Run 3) | $ELP_{S18}$ | 573 |
| | 2026.02.21 (Run 2) | $H5_{2/3}$ | 429 |
| | 2026.02.06 (Run 1) | H5 | 570 |
| | 2026.02.21 (Run 4) | H5-4Pepper | 758 |
| | 2026.02.21 (Run 3) | H5-H5 | 900 |

Transcription reactions contained 40 mM HEPES (pH 7.4), 80 mM KCl, 8 mM $MgCl_2$, 6 mM total NTP, 76 nM linear DNA, 50 nM T7 RNAP, and 10 mg mL$^{-1}$ GFP(+4).

**Table 6** Primers for Generating Linear DNA Templates for IVT

| Plasmid Template | Forward Primer | Reverse Primer | DNA Length (bp) |
|---|---|---|---|
| pET-6xHis-GFP(0) | T7 Promoter[1] | AGCGCCACCGTGATGGT | 120 |
| pET-6xHis-GFP(0) | T7 Promoter[1] | TGTTGTGCAAATAAACTTCA | 267 |
| pET-6xHis-GFP(0) | T7 Promoter[1] | GTTTCATATGATCAGGGTAACGACTGAAGCACTGAACG | 355 |
| pET-6xHis-GFP(0) | T7 Promoter[1] | TTATAACGCAGTTTATGGCCTAAAATGTTG | 548 |
| pET-6xHis-GFP(0) | T7 Promoter[1] | T7 Terminator[2] | 1002 |
| pET-6xHis-GFP(0)-4Pepper | T7 Promoter[1] | T7 Terminator[2] | 1157 |
| pET24a(+)-6xHis-$ELP_{S6}$ | T7 Promoter[1] | T7 Terminator[2] | 410 |
| pET24a(+)-6xHis-$ELPS_{S18}$ | T7 Promoter[1] | T7 Terminator[2] | 590 |
| pET24a(+)-6xHis-$H5_{2/3}$ | T7 Promoter[1] | T7 Terminator[2] | 446 |
| pET24a(+)-6xHis-H5 | T7 Promoter[1] | T7 Terminator[2] | 587 |
| pET24a(+)-6xHis-H5-4Pepper | T7 Promoter[1] | T7 Terminator[2] | 775 |
| pET24a(+)-6xHis-H5-H5 | T7 Promoter[1] | T7 Terminator[2] | 917 |

[1] TAATACGACTCACTATAGGG

[2] CAAAAAACCCCTCAAGACC

**Table 7** Key Resources Table

| Reagent or Resource | Source | Identifier |
|---|---|---|
| **Baterial Strains** | | |
| NEB 5-alpha Competent *E. coli* (High Efficiency) | New England Biolabs | Cat#C2987H |
| NiCo21(DE3) Competent *E. coli* | New England Biolabs | Cat#C2529H |
| **Chemicals, Peptides, and Recombinant Proteins** | | |
| Yeast Extract | Alfa Aesar | Cat#J23547-A1 |
| Enzymatic Digest Casein Hydrolysate | Thermo Fisher Scientific | Cat#AAJ12855P5 |
| Sodium Chloride | Thermo Fisher Scientific | Cat#S2713 |
| Bacto Agar | Becton Dickinson | Cat#214010 |
| Ampicillin (Sodium) | GoldBio | Cat#A-301-5 |
| QIAprep Spin Miniprep Kit | Qiagen | Cat#27106 |
| Phusion High-Fidelity DNA Polymerase | New England Biolabs | Cat#0530L |
| DpnI | New England Biolabs | Cat#R0176S |
| TopVision Agarose | Thermo Fisher Scientific | Cat#R0492 |
| QIAquick PCR Purification Kit | Qiagen | Cat#28106 |
| NEBuilder HiFi DNA Assembly Master Mix | New England Biolabs | Cat#E2621L |
| HiScribe T7 Quick High Yield RNA Synthesis Kit | New England Biolabs | Cat#E2050S |
| RNase R | New England Biolabs | Cat#M0100S |
| ssRNA Ladder | New England Biolabs | Cat#N0362S |
| Ribonucleotide Solution Set | New England Biolabs | Cat#N0450S |
| Monarch Spin RNA Cleanup Kit | New England Biolabs | Cat#T2050S |
| Isopropyl-beta-D-thiogalactoside (IPTG) | GoldBio | Cat#I2481C |
| HisPur Ni-NTA Resin | Thermo Fisher Scientific | Cat#88221 |
| Sodium Phosphate, Monobasic | Thermo Fisher Scientific | Cat#AC448170010 |
| Imidazole | Sigma-Aldrich | Cat#I202 |
| HEPES | Sigma-Aldrich | Cat#H3375-500G |
| Potassium Chloride | Sigma-Aldrich | Cat#P9541-500G |
| Magnesium Chloride Hexahydrate | Thermo Fisher Scientific | Cat#M33-500 |
| TBE Running Buffer (5X) | Invitrogen | Cat#LC6675 |
| MOPS-SDS Running Buffer (20X) | Thermo Fisher Scientific | Cat#J62847.AP |
| NuPAGE LDS Sample Buffer (4X) | Thermo Fisher Scientific | Cat#NP0007 |
| mPEG-Silane, MW 5000 | Laysan Bio | Cat#MPEGSIL50001G |
| Quick Start Bradford Protein Assay | Bio-Rad | Cat#5000201 |
| SYBR Green II RNA Gel Stain (10,000X) | Invitrogen | Cat#S7564 |
| SYTO 63 Nucleic Acid Stain | Invitrogen | Cat#S11345 |
| Alexa Fluor 594 NHS Ester | Invitrogen | Cat#A20104 |
| Cy5-UTP | Apexbio Technology | Cat#B8333 |
| DAPI | Sigma-Aldrich | Cat#D9542-1MG |
| **Deposited Data** | | |
| Microscopy Data and Segmented Masks | This paper | Figshare (doi.org/10.6084/m9.figshare.32136637) |
| **Oligonucleotides** | | |
| ssDNA Oligos | IDT | See Table 6 |
| **Recombinant DNA** | | |
| p6XHis-T7(P266L) | Chillon et al. | Addgene#174866 |
| pET-6xHis-sfGFP | Yeong et al. | Addgene#199171 |
| pET-6xHis-GFP(0) | Yeong et al. | Addgene#199166 |
| pET-6xHis-GFP(+4) | This paper | Addgene#256589 |
| **Software and Algorithms** | | |
| FIJI ImageJ (version 2.16.0/1.54p) | NIH | imagej.net/software/fiji |
| Cellpose-SAM (2025) | Pachitariu et al. | github.com/MouseLand/cellpose |
| Prism (version 10.6.1) | GraphPad | graphpad.com |
| Adobe Illustrator (2025) | Adobe | adobe.com |
| **Other** | | |
| 384-well Glass-Bottom Plate | Cellvis | Cat#P384-1.5H-N |
